# Geometric determinants of time series learning and attractor stability in closed-loop circuits

**DOI:** 10.64898/2026.11.10.765266

**Authors:** Alessandro Barri, Harsha Gurnani, R. Angus Silver

## Abstract

Skilled volitional behaviours arise from recurrent circuits, which can sustain structured neural activity in the absence of external drive, through internal feedback. But how such autonomous activity is learned and stability maintained in distributed motor circuits is poorly understood. To investigate the relationship between neural representations and learning we compared a cortex-like recurrent neural network (RNN) and a cortico-cerebellar-like circuit with a feed-forward layer (RNN+FFL), both in a closed-loop configuration. RNN+FFL circuits learned periodic target signals that parameter-matched RNNs could not, due to a failure to form attractor states. Decomposing population activity by frequency, rather than variance, separated the task-relevant and task-irrelevant subspaces. Quantifying their geometric arrangement revealed that the feed-forward layer improved learning and stabilised attractor states by increasing the angles within and between these subspaces. Our results show that task-specific representational geometry is a better predictor of periodic time series learning in recurrent circuits than classical global measures such as dimensionality.

## Introduction

Physically engaging with our environment requires temporally precise and coordinated muscle activation. The nervous systems of animals must therefore generate sequences of motor commands^1–3^ that drive actions ranging from locomotion^4^ and limb movement^5^ to whisking^6^, birdsong^7^ and speech^8^. Many of these time-varying signals, or time series, are learned and subsequently generated internally, allowing animals to generate volitional movements^9^. Such internally generated (autonomous) signals are widely thought to arise from recurrent dynamics, where internal feedback sustains structured activity in the absence of external drive^10–12^. But how circuits with multiple recurrent loops and nonlinearities learn and stably maintain the neural dynamics underlying the complex time series required for skilled movements remains poorly understood.

The motor cortex plays a key role in motor control and learning, generating the sequences of motor commands for volitional movements^3,10,13^, although its necessity for executing highly practiced, automatic sequences depends on task demands^14^. It is highly interconnected and forms both local and extended loops incorporating other brain regions. The largest of these is the cortico-cerebellar system^15,16^, which is critical for sensorimotor learning and skilled execution^17,18^. Intriguingly, this closed-loop combines distinct recurrent and predominantly feed-forward circuit architectures of the motor cortex and cerebellar cortex, respectively. Sensory and motor cortex project to the cerebellum via the basal pontine nucleus, where sensory information and efference copies of motor commands reach cerebellar granule cells through mossy fibres^19^. Divergence onto the vastly more numerous granule cells expands mossy fibre activity into a high-dimensional representation^20^, before it converges onto Purkinje cells (**Fig. 1a**). Purkinje cells innervate the cerebellar nuclei, whose output returns to the same cortical regions via thalamus, closing the loop^15,21^. If local recurrent circuits in the motor cortex are sufficient to generate time series autonomously this anatomical arrangement seems puzzling. What advantage is gained by adding a feed-forward layer in a closed-loop formation?

**Figure 1.**
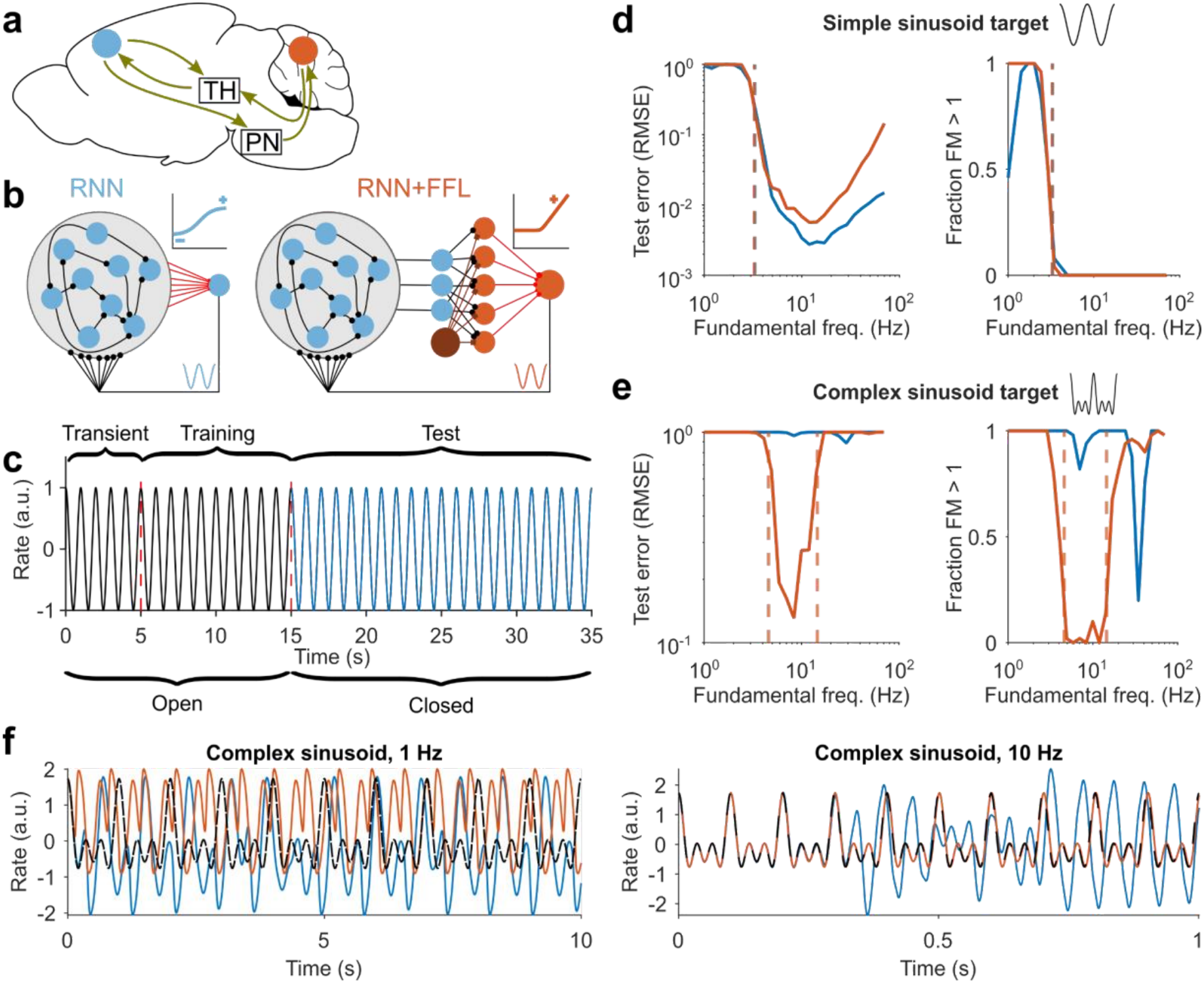
Addition of a cerebellar-like feedforward layer improves time series learning performance and stability in a cortex-like closed-loop recurrent neural network. **a** Schematic of the cortico-cerebellar system. Blue circle: motor cortex. Orange circle: cerebellar cortex. TH: Thalamus. PN: Pontine nucleus. Mouse brain outline adapted from Jonas Töle (Wikimedia Commons, CC0). **b** Left: scheme of simplified recurrent neural network (RNN) cortical model with tunable connections (red) to a readout neuron; inset: RNN hyperbolic tangent activation function. Right: scheme of simplified cortico-cerebellar model with cerebellar-like feed-forward layer and RNN (RNN+FFL); brown circle: inhibitory Golgi cell; inset: feed-forward layer rectified linear activation function. Tunable weights are shown in red. Example of target signal (blue/orange) shown on feedback branch. **c** Illustration of learning procedure. During the transient and training phases the behavioural task-relevant target signal is fed into the RNN (open loop, black line); weights are adjusted based on network activity during the training phase. During the test phase the RNN receives feedback from the readout unit (closed-loop, blue line). **d** Learning performance for a single cosine signal training. Left: Test error expressed as normalised Root Mean Square Error (RMSE) versus target signal fundamental frequency; lines represent averages over 50 simulations; blue line: RNN only; orange line: RNN+FFL; vertical dashed line: stability margin obtained from largest non-trivial Floquet multiplier; the coding level of units in the feed-forward layer was 0.5. Right: fraction of largest non-trivial Floquet multipliers larger than 1 versus signal fundamental frequency. **e** Same as (**d**), but for complex signal with 3 periodic components (Methods). **f** Sample complex signal outputs for two different fundamental frequencies. Dashed black line: target signal, RNN blue and RNN+FFL orange.

Autonomous time series learning has been studied mainly in recurrent neural networks (RNNs) with explicit feedback trained under the reservoir computing framework^22–24^. Recent analysis of a linear RNN showed that the dimensionality of the neural representation predicts learning performance for a single sinusoidal target^25^. This result might explain the involvement of the cerebellum in time series learning, given its proposed role in increasing the embedding dimensionality of neural representations through expansion recoding^26–28^. However, recent experimental measurements from the cortico-cerebellar system show that population activity in layer 5b of motor cortex and in cerebellar granule cells is highly correlated^18^. Moreover, cerebellar feedback through thalamus is necessary for maintaining preparatory activity in the motor cortex during goal-directed behaviours^29,30^ and for context-dependent movement initiation^21^. These findings have motivated proposals that cerebellar feedback stabilises cortical activity, corrects deviations^31,32^, and governs transitions between cortical attractor states^33^. This aligns well with the view that motor execution is controlled by sequences of attractor states^34^, and that periodic attractor dynamics underlie cyclic components of motor commands^35^. Consistent with this, the cortex and cerebellum communicate via distinct frequency bands^36^ and the cerebellum can maintain the frequency representations required for periodic movements^37^. While recent modelling of the cortico-cerebellar system has shown that the cerebellum can support cortical learning by providing context separation^38^, predictive feedback^39^ or fast error correction^40^ and facilitate task acquisition and consolidation^41^, these studies have not explored how neural representations within cortico-cerebellar loops influence time series learning or the stability of attractors underlying rhythmic movements, which remains poorly understood.

To identify the fundamental principles of autonomous times series learning in recurrent motor systems, we studied reduced, rate-based models as these facilitate numerical simulations and conceptual interpretation. We compared the neural representation and learning performance of cortical-like closed-loop RNN circuits to networks combining an RNN with a cerebellar-like feed-forward layer (RNN+FFL) with an identical number of plastic readout weights (parameter-matched). RNN+FFL circuits outperformed RNN-only circuits, when trained to autonomously generate the complex time series required for periodic motor tasks. To understand why, we developed a Fourier-based dimensionality reduction approach that decomposes population activity by frequency, since application of conventional principal component analysis (PCA) failed to provide clear insights. Our Fourier decomposition revealed that the feed-forward layer separates task-relevant frequency components required for learning from undesirable, task-irrelevant components. Frequency separation, which was increased by inhibition and neuronal heterogeneity, reduced interference between task-relevant and irrelevant components thereby improving the stability of periodic attractor states. We establish the generality of the mechanism by showing that linear networks reproduce the stability properties of non-linear networks once their representational geometry is qualitatively matched. Our framework for quantifying the angular separation and amplitude of frequency components shows that task-specific representational geometry is more informative than global dimensionality for understanding time series learning and attractor stability in closed-loop network architectures.

## Results

### A cerebellum-like feed-forward layer extends the range of frequencies over which a recurrent neural network can learn complex periodic time series

To capture the essence of the cortico-cerebellar loop circuitry (**Fig. 1a**), while still making investigation tractable, we assumed that interconnecting structures between the motor cortex and the cerebellar cortex, act as relays. Following previous work we modelled the recurrent connectivity of the motor cortex as an RNN of N nonlinear units, which projected plastic synapses to a readout unit that fed back onto the network in a closed-loop configuration^22,25^ (**Fig. 1b**; **Methods**). The largely feedforward architecture of the cerebellar cortex was modelled as a two-layer perceptron^26,27^ in which mossy fibres relayed cortical activity to a feed-forward layer of instantaneous granule cell-like units with a smooth rectification under global Golgi cell inhibition, operating in a fluctuation-driven regime^42,43^, with a single linear Purkinje-like readout returning its output to the RNN (**Fig. 1b**; **Methods**). Only granule-cell-to-Purkinje-cell weights were plastic. We refer to this circuit as RNN+FFL and compare it to the parameter-matched closed-loop RNN. As our aim was to study the impact of representational geometry on learning performance, rather than the temporal dynamics of learning, we fixed the recurrent weights within the RNN. Likewise, we trained the readout weights of both models with least-squares regression to minimise dynamics introduced by training optimisation.

To evaluate the performance of the two models we trained the read-out unit to autonomously generate target signals with different numbers of periodic components. Training involved driving the circuit with the target signal in an open loop until activity reached steady state, fitting readout weights using least squares, and assessing performance during a closed-loop test phase using the root-mean-square error between generated and target signals (**Fig. 1c**). For target signals with a single periodic component (cosine), the two circuits performed comparably across a range of fundamental frequencies (**Fig. 1d**, left; n = 50 networks per condition**; Fig. S1** for individual errors for each simulation). Both showed high error at low frequencies, a minimum at intermediate frequencies, consistent with previous work^25^, and a degradation in performance thereafter. Because a failure to learn the periodic target signal could reflect either the failure of the network to form a periodic attractor, or the formation of an attractor that was inaccurate or exhibited unwanted network dynamics, we applied Floquet analysis^44–46^ while keeping the dynamics on the target trajectory (**Methods**). This enabled us to determine whether the feedback destabilised the desired periodic attractor after training. Calculation of the largest non-trivial Floquet multiplier of the target periodic solution, revealed values greater than unity in the high-error region, indicating that the failure of the circuit to learn arose from an inability to form the desired stable periodic attractor at low frequencies (**Fig. 1d, right; Fig. S1** for individual Floquet multipliers for each simulation).

Complex movements require motor commands containing multiple frequency components. To investigate whether RNN and RNN+FFL circuits could learn more complex autonomous signals we trained both circuits on target signals with multiple frequency components. The RNN failed at every fundamental frequency tested, whereas the RNN+FFL reproduced the signal with low errorc for fundamental frequencies between 5 and 10 Hz (**Fig. 1e**,**f; Fig. S1**). Floquet analysis predicted this difference in performance, with multipliers exceeding unity almost everywhere across the full range for the RNN, but remaining below unity throughout the RNN+FFL’s low-error band (**Fig. 1e, right; Fig. S1**). These results suggest that adding a cerebellum-like feed-forward circuit to a motor-cortex-like recurrent circuit, in a closed-loop formation, improves autonomous time series learning by enabling the formation of stable periodic attractors that encode multi-frequency signals.

### Target-signal-matched neural dynamics revealed with Fourier-based decomposition

Why does a feed-forward layer enable the formation of stable periodic attractor states? To investigate this question we compared the population activity in the two circuits, since any differences in representations could influence how effectively the readout synapses learn the target signal.

We first examined a simple linear RNN in open loop configuration driven by a single frequency target signal, since this case is well understood^25^ (**Fig. 2a**). The activation rates of the driven RNN units were phase-shifted versions of the input signal (**Fig. 2b**). As has been shown analytically, this is equivalent to the collective dynamics being embedded in a 2-dimensional plane within the RNN’s N-dimensional state space^25^. Consistent with this, Principal Component Analysis (PCA) of RNN activity revealed the existence of two modes that had identical frequencies, but were phase shifted with respect to the target signal (**Fig. 2c**). By contrast, when we applied a Fourier decomposition to the network’s activity, one of the components was precisely aligned to the target signal (magenta line, **Fig. 2d**). Fourier analysis therefore isolates the periodic activity mode (i.e. the cosine component) to be learned.

**Figure 2.**
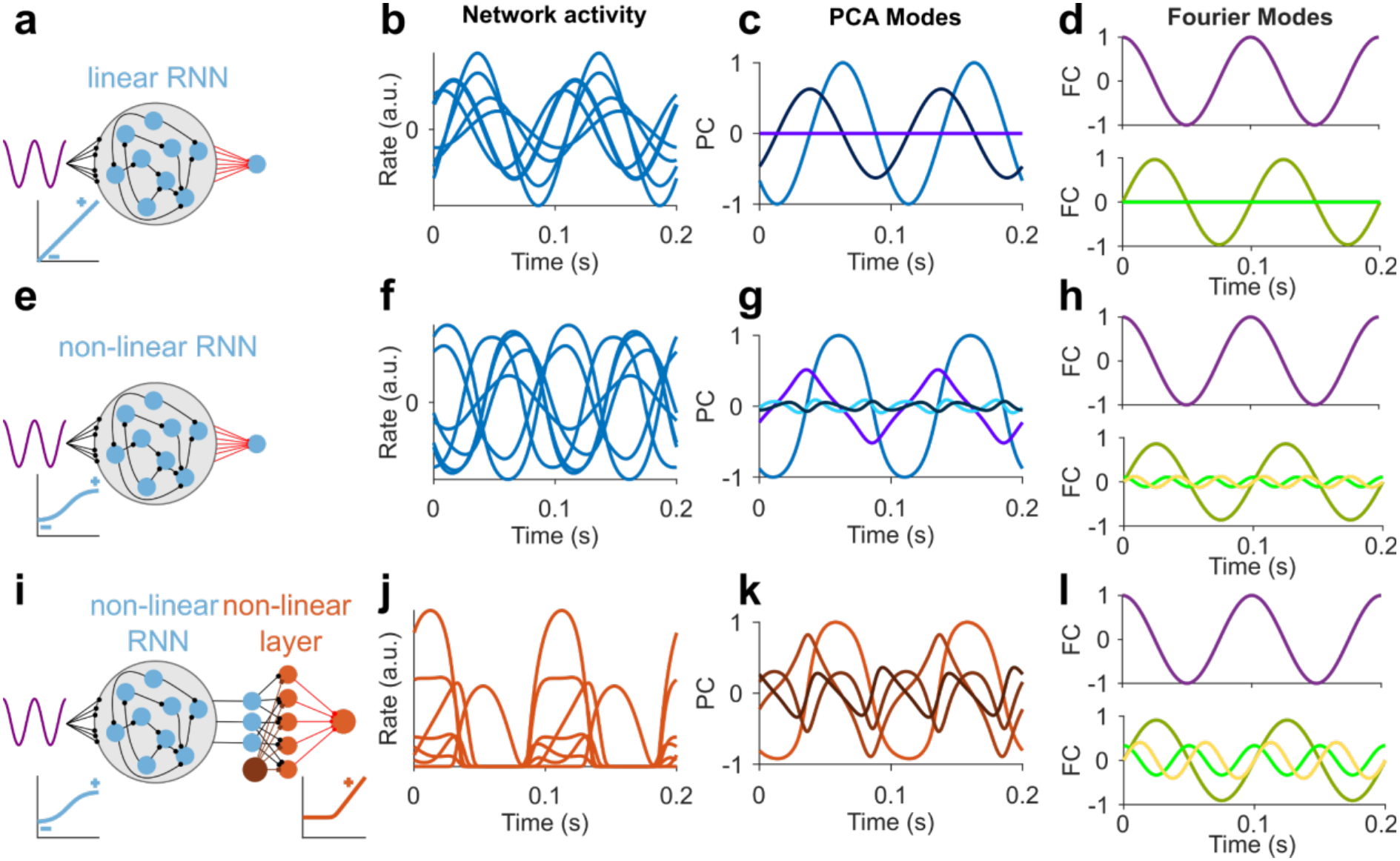
Fourier decomposition reveals that nonlinear neuronal activation functions introduce additional frequency components consisting of signal harmonics. **a** Schematic of cortical-like recurrent neural network (RNN) with linear activation function driven by simple cosine target signal. **b** Sample population activity of the linear RNN in response to cosine signal input. Each line represents the activity of a single unit. **c** First four principal component modes of population activity. Only the first two modes are non-zero. **d** First four Fourier modes of the RNN activity. Upper panel: magenta line denoting task-relevant vector mode. Lower panel: green and yellow lines denote task-irrelevant vector modes; only one non-zero mode is present. **e-h** Same as (**a-d**) but for nonlinear RNN with tanh activation function. For clarity, in (**h**) only non-zero modes are shown. **i-l** Same as (**a-d**) but for nonlinear cortico-cerebellar RNN+FFL with non-linear units in both the RNN and feed-forward layer. PCA and Fourier modes are for feed-forward cell layer activity.

For non-linear RNNs (**Fig. 2e**) individual activation rates were distorted versions of the target signal, an effect that arose from the nonlinear activation functions (cf. **Fig. 2f and 2b**). Unlike the linear case, trajectories left the two-dimensional plane, and every principal component contained multiple frequencies, making them difficult to relate to the target signal (**Fig. 2g**). Fourier decomposition naturally separated neural population activity into the target signal (**Fig. 2h**, magenta line), and non-target signal components with either different frequencies and/or phases (**Fig. 2h**, green and yellow lines). Re-coding through the feed-forward layer amplified this effect of nonlinear units. Activations of feed-forward layer units were more distorted than RNN activations (**Fig. 2i**,**j**), and both higher-order principal components and higher-frequency Fourier modes carried more variance (**Fig. 2k**,**l**). This analysis shows that neuronal non-linearities introduce additional frequency components (e.g. harmonics) that could potentially interfere with the network’s ability to learn the target signal.

A similar picture emerged when we trained networks on more complex target signals, composed of 3 frequency components (**Fig. 3a**). The activity of individual units and principal components in the RNN contained multiple frequency components (**Fig. 3b,c**), whereas Fourier decomposition recovered three activity modes in the RNN matching the frequencies and phases of the target signal harmonics (**Fig. 3d**). Interestingly, the RNN+FFL represented the same three signal components alongside a larger set of non-target-signal components (**Fig. 3e-h**). Despite this, the performance of the nonlinear RNN+FFL was superior to the RNN when learning complex signals (**Fig. 1e**).

**Figure 3.**
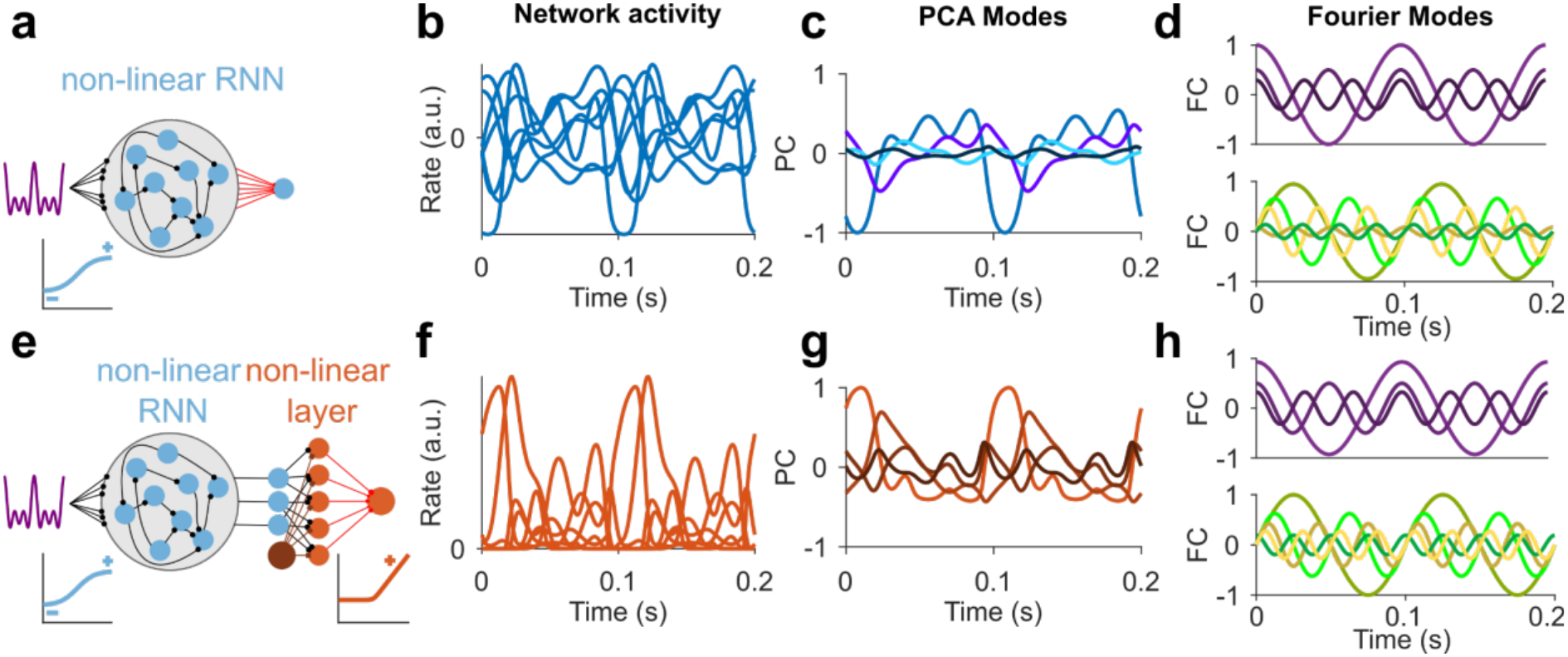
Fourier decomposition of complex target signal representations separates task-relevant and task-irrelevant activity modes in non-linear RNN and RNN+FFL. **a** Schematic of recurrent neural network (RNN) with non-linear activation function driven by complex signal. **b** Sample population activity of the non-linear RNN in response to complex signal input. Each line represents the activity of a single unit. **c** First four principal component modes of the population activity. **d** First eight Fourier modes of the population activity. Upper panel: magenta lines denote task-relevant vector modes. Lower panel: green and yellow lines denote task-irrelevant vector modes with higher frequencies. **e-h** Same as (**a-d**) but for nonlinear cortico-cerebellar RNN+FFL circuit.

Our Fourier-based approach decomposes network activity into two types of frequency components: those that encode the *task-relevant* target frequencies and ‘noisy’ *task-irrelevant* activity modes that arise from RNN dynamics and neuronal nonlinearities. Fourier decomposition therefore provides clear advantages over PCA for studying periodic time series learning in both linear and nonlinear networks. It is clear from our analysis that the internal representations in linear and nonlinear networks differ, and our naive expectation was that the generation of ‘noisy’, task-irrelevant components by nonlinear units should impair learning performance rather than improve it. Moreover, the increased presence of task-irrelevant components in nonlinear RNN+FFLs is also at odds with their superior performance over nonlinear RNNs in learning and stably generating complex target signals (**Fig. 1e**).

### A geometric framework to link neural representations to closed-loop stability

To resolve this apparent paradox we investigated whether the difference in learning performance and stability in nonlinear RNN and RNN+FFL circuits arose from the geometrical arrangement of task-relevant and task-irrelevant components in neural activity state space. To link the geometry of the neural population activity to learning performance we treated each frequency component within the population activity as a vector, and asked how its geometric arrangement aligns with the vector of synaptic readout weights (**Fig. 4a**). Importantly, frequency components, unlike principal components, are not orthogonally arranged in state space.

**Figure 4.**
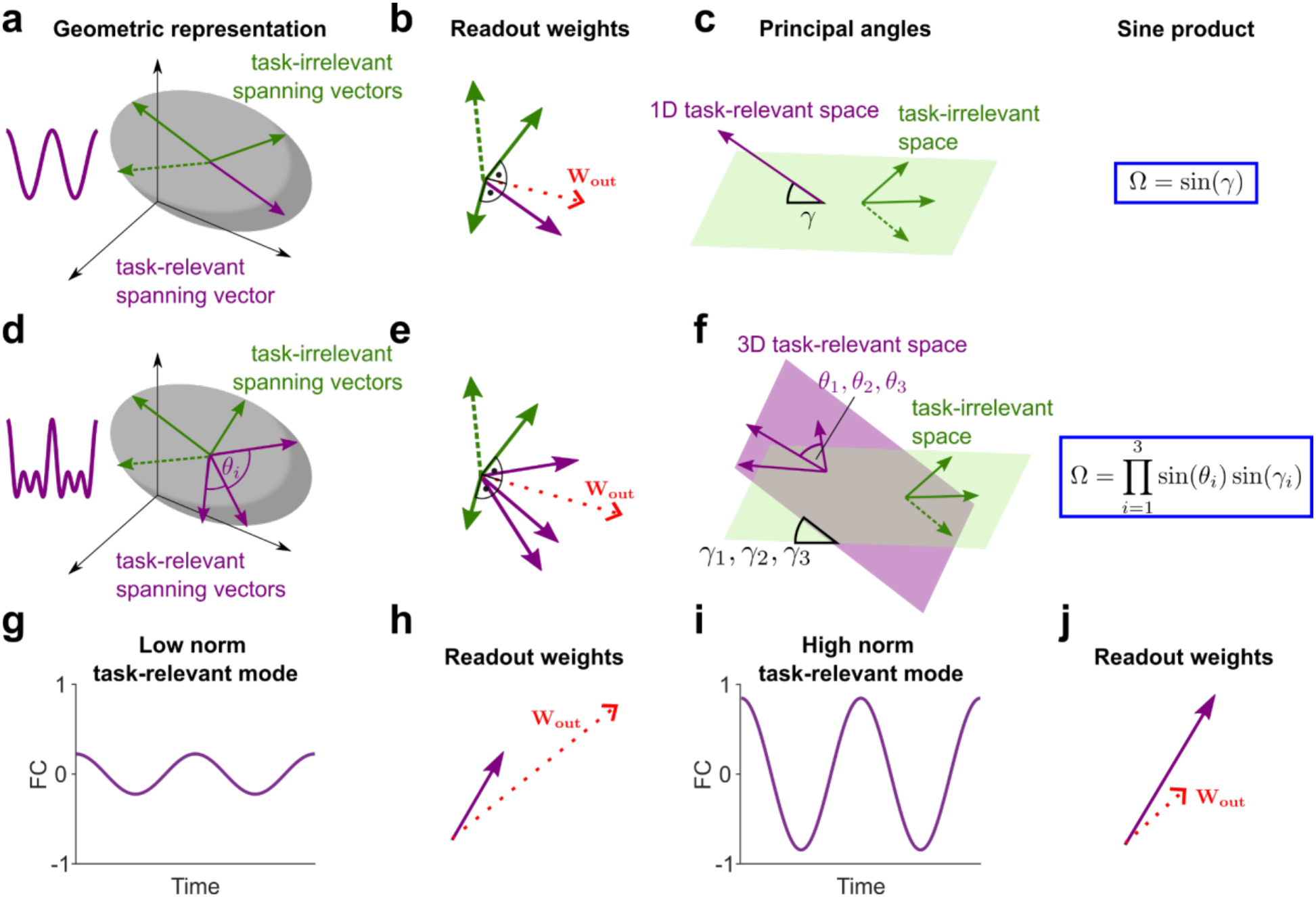
Geometric properties of neural representations in non-linear networks for simple and complex target signals. **a** Schematic view of population activity of non-linear network projected into low-dimensional subspace for cosine target signal showing task-relevant (magenta) and task-irrelevant (green) spanning vectors. **b** Illustration of readout weight vector *W*_*out*_ (red) and its relationship to task-relevant spanning vector (magenta arrow), and task-irrelevant spanning vectors (green arrows). **c** Left: Illustration of principal angle between task-relevant and task-irrelevant subspaces, denoted by *γ*. Right: Definition of the sine product for the simple target signal case. **d-f** Same as (**a-c)** but for complex target signal with three frequency components. The angles between task-relevant spanning vectors are denoted by *θ*_*i*_. **g** Example of simple signal mode (task-relevant fourier component, FC) with comparatively low amplitude. **h** Task-relevant spanning vector with low norm leading to readout vector with large norm. **i** Example of simple signal mode with comparatively high amplitude. **j** Task-relevant spanning vector with high norm leading to readout vector with low norm.

We hypothesised that three conditions should govern stable learning. First, the readout weight vector should align with the vector(s) encoding the target signal (task-relevant spanning vectors), while remaining orthogonal to vectors encoding task-irrelevant frequencies or phases, since any overlap between these will feed unwanted frequencies or phase-shifts back into the loop (**Fig. 4b,e**). In general, multiple task-relevant and multiple task-irrelevant components are present. Thus, we consider the two subspaces spanned by each of these sets, the task-relevant-subspace and task-irrelevant-subspace, respectively, and quantify their separation by calculating the principal angles between them. Intuitively, the principal angles generalise the notion of an angle between two vectors, to subspaces, as they capture how one subspace must be rotated to align with another (**Fig. 4c,f**; **Methods**). Second, for target signals with more than one frequency component, when there are several task-relevant spanning vectors with unequal amplitudes (norms), these vectors should be separated from one another (**Fig. 4e**). This is because closely aligned vectors cause the least-squares solution to be poorly conditioned, making it sensitive to noise (**Fig. 4f**), as we illustrate for the analytically tractable case for two task-relevant vectors and one task-irrelevant vector (**Fig. S2**; **Methods**). Third, we hypothesise that representations with large task-relevant vector amplitudes should be more stable than those with low amplitudes, since the norms of the readout weight vector and the task-relevant vectors are inversely related (**Fig. 4g-j**), and large readout norms are associated with unstable feedback dynamics^25,47^.

To measure whether these conditions lead to stable time series learning we developed two metrics. The first two conditions involve geometric separation between frequency components and we combine them by calculating a single quantity, the product of the sines of the angles between task-relevant vectors and the principal angles (**Fig. 4c,f**; **Methods**). This *sine product* approaches 1 when all components are mutually orthogonal and collapses to zero when either the task-relevant components align with each other or the task-relevant and task-irrelevant subspaces align in at least one direction. The third condition quantifies the strength of the signal. We assay it by calculating the average norm of the task-relevant vectors. These measures provide a framework to link the geometry of neural representations to learning performance and stability in closed-loop neural circuits.

### Testing predicted geometric determinants of time series learning by mimicking nonlinear behaviour in tractable linear circuits

Testing whether geometric parameters of neural state space predict closed-loop stability and learning performance is challenging in nonlinear circuits because the angles and amplitudes of task-relevant and task-irrelevant vectors covary in complex ways when a single variable is altered. To disentangle the effects of altering principal angles from task-relevant vector norms, we designed a tractable linear RNN+FFL network that qualitatively mimicked the task-irrelevant components of nonlinear networks by injecting artificial activity into the feed-forward layer that resembled the task-irrelevant components in nonlinear networks. (**Methods**; **Fig. 5a,b**). Without this task-irrelevant input the frequency components in the linear feed-forward layer are identical to the components generated by the linear RNN (i.e. cosine signal and non-signal sine; **Fig. 2d**). By contrast, when the artificial activity was introduced an additional task-irrelevant component was present in the feed-forward layer **(Fig. 5b**, green arrow). Adjustment of the correlation between the synaptic weights of the artificial input and the cosine task-relevant-spanning vector in the feed-forward layer controlled the angle between these two components (α). This enabled us to continuously manipulate the principal angle between the task-relevant and task-irrelevant subspace (*γ*), when multiple task-irrelevant components were present, while keeping all the spanning vector norms fixed (**Fig. 5b; see Methods**).

**Figure 5.**
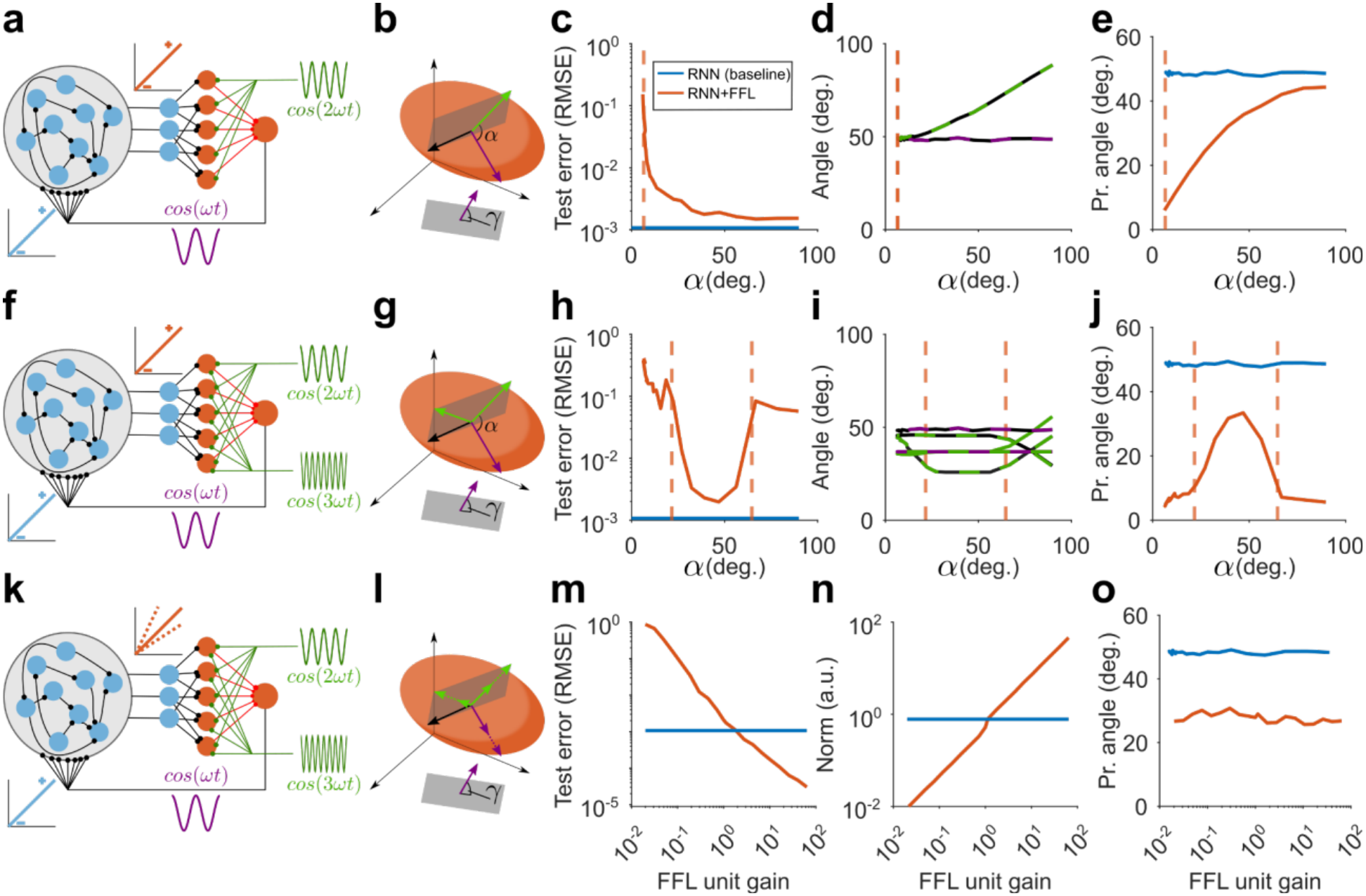
The principal angle predicts collapse of closed-loop stability in linear RNN+FFL circuits with artificial task-irrelevant input. **a** Scheme of linear RNN+FFL with artificial task-irrelevant input (green) injected into the feed-forward layer. **b** Schematic view of resulting population activity projected into low-dimensional subspace. Magenta arrow: task-relevant spanning vector; black arrow: task-irrelevant spanning vector inherited from RNN dynamics; green vector: artificially injected task-irrelevant spanning vector; grey plane: task-irrelevant-subspace; α: adjustable angle between task-relevant and artificial task-irrelevant vector; *γ* : principal angle between task-relevant and task-irrelevant subspaces. **c** Average test error (root mean square error; RMSE) versus α; lines are averages over 50 simulations. Orange line: RNN+FFL; blue line: baseline performance of RNN only; dashed vertical orange line: stability margin obtained from largest non-trivial Floquet multiplier. **d** Individual angles between spanning vectors versus α; lines correspond to the angle between the two task-irrelevant spanning vectors (black-green line; black and green arrows in (**b**)) as well as the angle between the task-relevant and RNN generated task-irrelevant spanning vectors (black-magenta line; magenta and black arrows in (**b**)). **e** Principal angle (*γ*) versus α (lines as for **c**) **f-j** As (**a-e**) but for two artificial task-irrelevant inputs. **i** Individual angles between spanning vectors with colour scheme analogous to the one in (**d**). **k-l** As (**f-g**) but for varying feed-forward layer gain and fixed angles resulting in *γ* ≈ 28 degrees. **l** Solid/dotted arrows represent task-relevant and task-irrelevant vectors affected by feed-forward layer unit gain changes. **m** Average test error versus feed-forward layer unit gain. **n** Task-relevant spanning vector amplitude (Norm) versus feed-forward layer unit gain. **o** Principal angle (*γ*) versus feed-forward layer gain.

As the angle between task-relevant and task-irrelevant spanning vectors was decreased from orthogonal, the closed-loop test error monotonically increased for a single cosine target signal (**Fig. 5c**). When the angle became small (α < 7 degrees), some simulations became unstable, as indicated by the appearance of Floquet multipliers that exceeded unity (**Fig. 5c**, dashed line). We confirmed that the angle between the task-relevant and task-irrelevant components in the RNN were unaffected by the artificial input to the feed-forward layer (black-magenta line, **Fig. 5d**). Thus, the fall in the principal angle between the task-relevant and task-irrelevant subspace (and thus the sine product), was directly attributable to manipulating α (**Fig. 5e**). Consistent with our framework, these simulations show that the value of the principal angle is inversely related to the closed-loop error and that instabilities appear when it becomes small.

To examine the impact of higher order task-irrelevant harmonics in non-linear networks (**Fig. 2i**) we injected a second, higher frequency artificial input component into the feed-forward layer (**Fig. 5f**) and adjusted the angle between the task-relevant component and the lower frequency artificial task-irrelevant component to manipulate the principal angle (**Fig. 5g**). Surprisingly, under these conditions the closed-loop test error was a non-monotonic function of α, with a region of comparatively good performance for intermediate values, with instabilities appearing at high and low values (**Fig. 5h**). While the angles between individual spanning vectors were heterogeneous and difficult to interpret (**Fig. 5i**), the principal angle, predicted the network’s close-loop performance and the region over which it can form stable periodic attractors (**Fig. 5j**).

To test whether amplitude of the task-relevant vector impacted learning we varied the gain of the units in the feed-forward layer, while keeping all other parameters fixed (**Fig. 5k**). This altered the task-relevant spanning vector norm without changing the principal angle between task-relevant and task-irrelevant components (**Fig. 5l**). As predicted, we found that the closed-loop test error decreased with increasing gain (**Fig. 5m**), due to the task-relevant vector norm increasing (**Fig. 5n**). Importantly, the principal angle remained unaffected by this manipulation (**Fig. 5o**).

These simulations show that the neural representations of non-linear closed-loop networks can be mimicked in more tractable, linear systems, by introducing artificial input harmonics. By selectively manipulating key geometric properties with this new approach, we confirmed our predictions that separating task-relevant and task-irrelevant components is critical for forming the periodic attractors required for stable closed-loop time series learning, with the principal angle (or sine product) serving as a key metric of this separation. Furthermore, we show that closed-loop performance is directly linked to the strength of task-relevant frequency modes, as quantified by the task-relevant spanning vector norms.

### Dimensionality is a poor predictor of time series learning performance for complex signals

Neural population activity is often quantified in terms of embedding dimensionality and/or condition number^25,48,49^ and these have been shown to be predictive of learning performance in linear RNNs for single component periodic signals^25^. However, when we extended this linear analysis to more complex time series the relationships between learning performance and dimensionality or condition number became less clear (**Fig. S3**). To test their predictive power more directly we varied the amplitude of the artificial input spanning vector in our linear networks, since this should alter the dimensionality and condition number of the representation, but leave the principal angle unchanged (**Fig. 6a,b**). Interestingly, the test-error did not change when varying the task-irrelevant vector norm over four orders of magnitude (**Fig. 6c**). By contrast, the embedding dimensionality peaked and the condition number reached a minimum when the ratio between task-relevant and task-irrelevant vector norms was close to unity (**Fig. 6d**). The principal angle, as quantified by the sine product, was constant across all tested norm ratios, similar to the test-error (**Fig. 6e**). These results suggest that unlike the sine product, dimensionality and condition number, two common global measures of population activity, fail to predict the performance of time series learning in closed-loop networks, when multiple task-irrelevant components are present.

**Figure 6.**
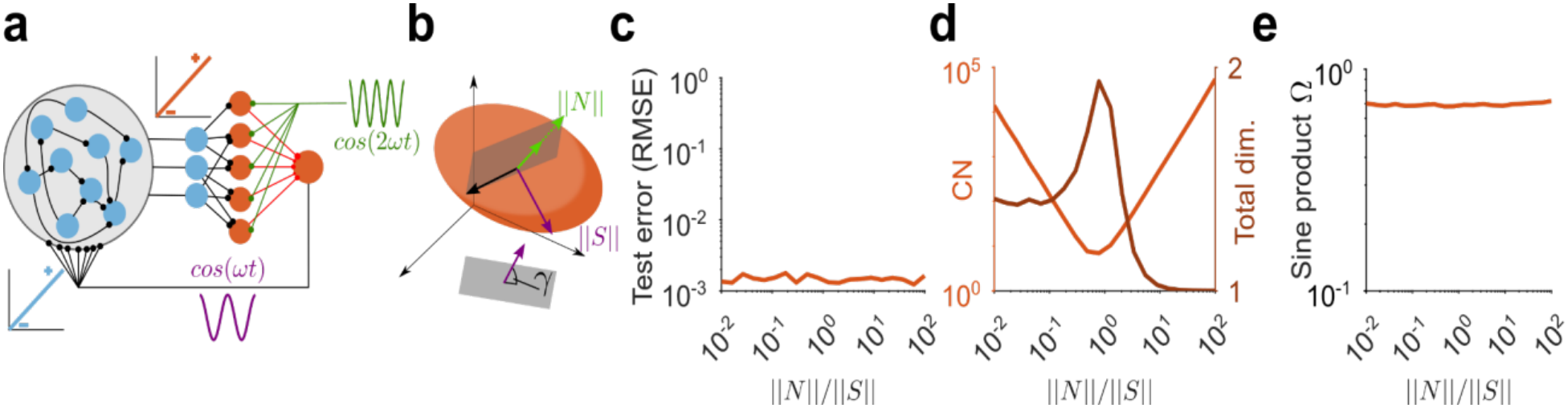
Dimensionality and condition number are not predictive of learning performance for linear RNN+FFL with artificial task-irrelevant input. **a** Schematic of linear RNN+FFL with artificial task-irrelevant input (green) injected into the feed-forward layer. **b** Schematic view of resulting population activity projected into low-dimensional subspace. ||*S*||: norm of task-relevant spanning vector (magenta arrow); ||*N*||: norm of injected task-irrelevant spanning vector (green arrow); *γ* : principal angle between task-relevant and task-irrelevant subspaces. **c** Average test error (root mean square error, RMSE) versus ratio of spanning vector norms; lines are averages over 50 simulations. **d** Condition-number (CN, light orange line) and total dimensionality (brown line) versus ratio of spanning vector norms. **e** Sine product versus ratio of spanning vector norms.

### Metrics of representational geometry predict learning and stability in non-linear cortico-cerebellar loops

We next tested whether the relationships between representational geometry, time series learning and closed-loop stability held for non-linear RNN+FFL networks operating at different activity levels. Activity was controlled by adjusting the threshold of nonlinear units in the feed-forward layer (**Fig. 7a**). For a single frequency target signal, learning performance decreased as the activity level was reduced and became unstable at 0.25 regardless of fundamental frequency. Consistent with our linear network simulations, we found that for each activity level the test error was lowest when the sine product was maximal (**Fig. 7b**). Although the sine product increased as input layer activity became sparser (**Fig. 7b**), its impact on learning performance was offset by a marked reduction in the amplitude of the task-relevant vector and the sharp fall-off in norms at higher fundamental frequencies (**Fig. 7c**). Importantly, at high levels of activity (low thresholds, fc=0.75), when the task-relevant vector norms of RNN and RNN+FFLs representation were very similar (**Fig. 7c**, blue and brown lines), the sine product was a good predictor of performance differences between RNN and RNN+FFL. These results suggest that as for the linear networks with artificial input, learning performance and network stability in nonlinear closed-loop networks is determined by a combination of the sine product and the task-relevant vector norm.

**Figure 7.**
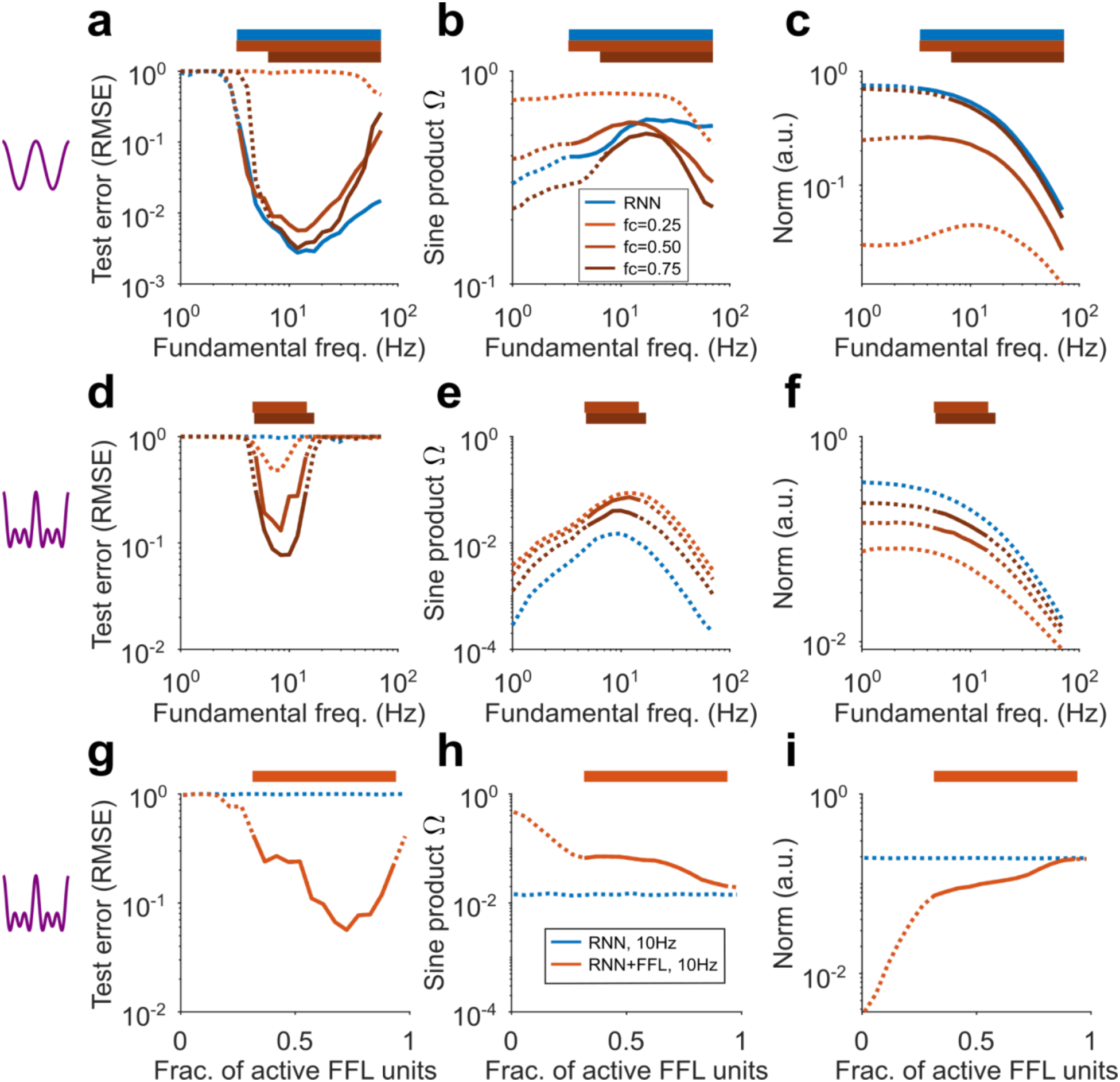
Geometric determinants of time series learning and stability in closed-loop non-linear networks across different levels of activity. **a** Test error versus fundamental frequency for simple target signal: nonlinear RNN (blue line) and nonlinear RNN+FFL (orange lines) at different activity levels in the feed-forward layer (fc, key in panel b); lines are averages over 50 simulations; horizontal bars and solid lines indicate regions of stability obtained via Floquet analysis; dashed lines indicate regions of instability. **b** Sine product (*Ω*) versus target signal fundamental frequency. **c** Amplitude of task-relevant subspace spanning vector (Norm) versus target signal fundamental frequency. **d-f** As (**a-c**) but for complex target signal. **g-i** Test error, sine product and average task-relevant vector norm versus fraction of active units in the feed-forward layer (orange line) for complex target signal at 10Hz; RNN shown for comparison (blue dashed line).

Nonlinear RNNs and RNN+FFLs exhibited narrower regions of stability for the complex target signal. Moreover, lowering activity in the feed-forward layer was associated with increased test errors (**Fig. 7d**). As for the single cosine case, the sine product maxima aligned approximately to the peak performance in the RNN+FFLs (**Fig. 7e**), but were smaller than for the single-frequency target, consistent with the overall drop in performance. However, RNN+FFL sine products were larger than the RNN sine product, regardless of the activity level (**Fig. 7e**). This, together with the moderate reduction of RNN+FFL task-relevant vector norms relative to the RNN (**Fig. 7f**) is consistent with the sine product and the task-relevant vector norm determining the learning performance and stability regions in the nonlinear RNN+FFL.

To investigate how learning performance and stability changes with feed-forward layer activity levels we examined the case for a complex signal at a fixed frequency (10 Hz). The RNN+FFL exhibited lower test errors than a parameter-matched RNN for all feed-forward layer activity levels larger than approximately 0.15 and had an optimum performance at approximately 0.75 (**Fig. 7g**). Floquet analysis identified a broad region of stability at intermediate activity levels (**Fig. 7g**). Consistent with this, the RNN+FFL sine product was strictly larger than the RNN sine product throughout (**Fig. 7h**). Importantly, the sine product monotonically increased with decreasing activity level (**Fig. 7h**), while the average task-relevant vector norm monotonically decreased with decreasing activity level (**Fig. 7i**) and exhibited a pronounced drop when instabilities appeared (**Fig. 7i**). This suggests that network stability was lost as a result of distinct failure modes: low signal strength (small norms) at low activity levels and low angular separation (low sine product) at high activity levels. Importantly, in the region of intermediate activity levels where the sine product was approximately constant, RNN+FFL performance was predicted by the average task-relevant vector norm (cf. **Fig. 7h and 7i**).

These simulations show that our framework for linking the geometry of neural representations to learning performance and stability extends to nonlinear closed-loop neural circuits. Our results suggest that network stability and task performance are maximised at intermediate activity levels in the feed-forward layer when the trade-off between the angular separation of frequency components and the amplitudes of the task-relevant components is optimal. These findings support our hypothesis that these two factors jointly predict closed-loop performance and stability at a qualitative level.

### Biologically inspired heterogeneity and inhibition improve time series learning

Our geometric framework for understanding closed-loop time series learning is based on homogeneous circuits and least-squares learning. To test whether its predictions hold under more realistic conditions we introduced cellular heterogeneity and a more biologically plausible learning rule. We employed a simplified FORCE variant^22^ incorporating the covariance rule^50,51^, in which synaptic weight updates are normalised by the norm of the population activity (**Methods**). This normalisation substantially reduces the influence of frequency-dependent task-relevant norms, so we assessed these networks using the sine product and stability alone. We compared homogeneous RNN and RNN+FFL circuits (**Fig. 8a**,**c-f**) with circuits in which neuronal time constants were heterogeneous (**Fig. 8b**,**g**). Heterogeneous RNN time constants were drawn from an exponential distribution matched in mean to the homogeneous case and those in the feed-forward layer were drawn from experimental measurements from granule cells^52^. We also examined the effect of Golgi cell feed-forward inhibition **(Fig. 8c**) and setting the ratio of feed-forward layer units to RNN units to 4:1 so as to approximately match the experimentally reported ratio between cortical and cerebellar neurons^53^ and the local mossy fibre to granule cell expansion ratio^18,54,55^ (**Fig. 8e-g**). White noise was injected into all units during the test phase (**Methods**).

**Figure 8.**
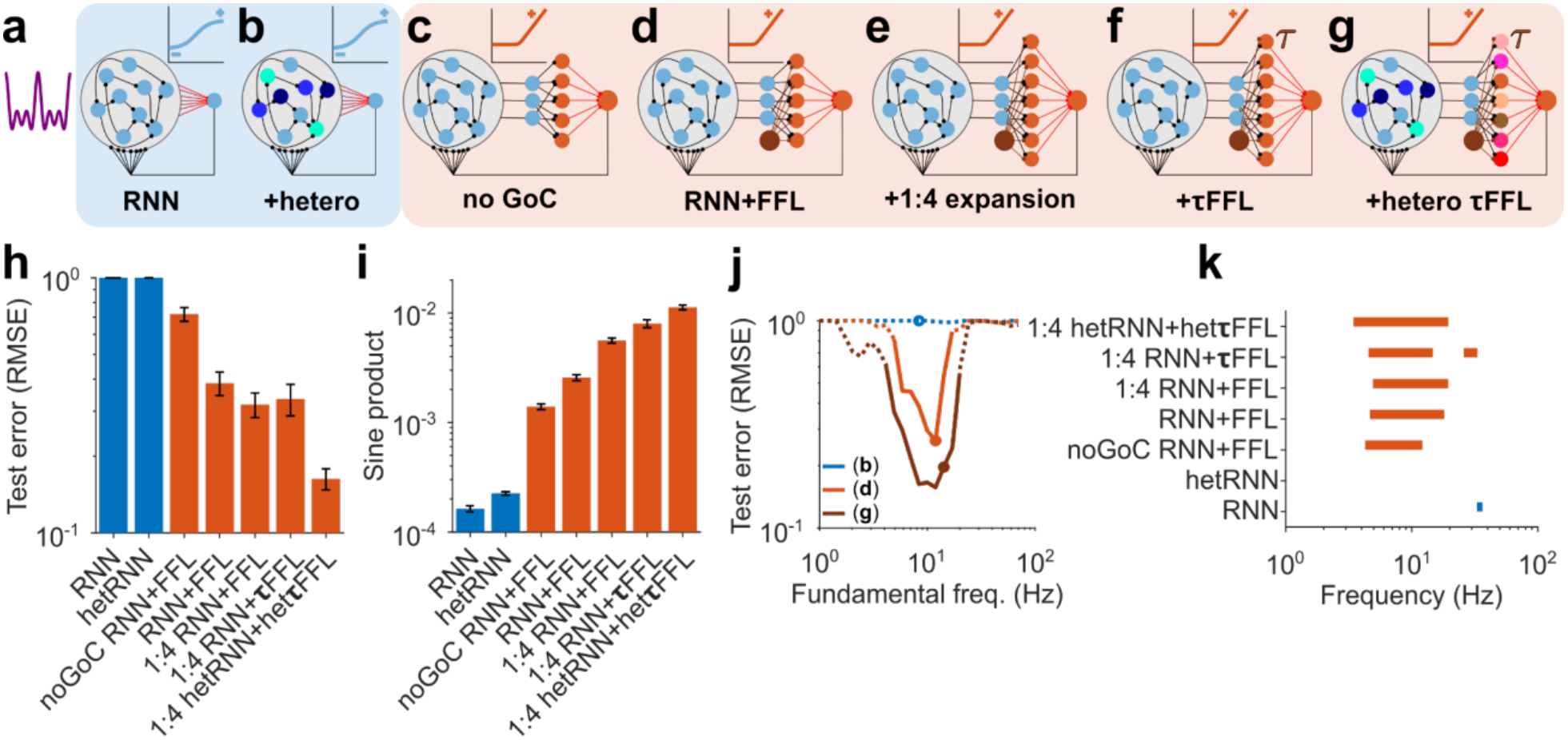
Biologically inspired neuronal heterogeneity improves time series learning performance and stability. **a-g** Schematics of different circuit configurations and complex signal to be learned. **a** Non-linear recurrent neural network (RNN) with homogeneous dynamical units. **b** Non-linear RNN with heterogeneous units (time-constants). **c** Nonlinear feed-forward layer with homogeneous instantaneous units plus RNN with homogeneous units (RNN+FFL), without Golgi cell (GoC) inhibition and 1:1 RNN unit to feed-forward layer expansion ratio. **d** As (**c**) but with Golgi cell inhibition. **e** As (**d**) but with 1:4 RNN unit to feed-forward layer expansion ratio. **f** As (**e**) but with homogeneous dynamical units in both the RNN and FFL (1:4RNN+***τ***FFL). **g** As (**f**) but with heterogeneous dynamical units (1:4hetRNN+het***τ***FFL). **h** Test error for different networks shown in (**a-g**) for a complex signal with fundamental frequency of 8.5Hz and feed-forward layer coding level of 0.4; bars are averages over 50 simulations for all networks except for (**f**) which was 24; error bars denote SEM. **i** Sine product *Ω* for different networks. **j** Test error versus target signal fundamental frequency for the networks from **b** (blue line), **d** (orange line) and **g** (dark orange line); dots indicate frequency for which the respective sine product is maximal. **k** Range of closed-loop stability across signal fundamental frequency for the different networks, measured by the largest non-trivial Floquet multiplier.

All circuits were trained on the complex target, for which the homogeneous nonlinear RNN is largely unstable. Heterogeneous time constants in the RNN alone did not reduce the test error, even at the near-optimal fundamental of 8.5 Hz (**Fig. 8a,b,h**). By contrast, a homogeneous RNN+FFL could learn the complex signal even without Golgi Cell feedforward inhibition, but the learning performance was substantially improved when Golgi cell inhibition was added (**Fig. 8c,d,h**). Increasing the number of cells in the feed-forward layer led to modest improvements in the test error (**Fig. 8d,h**). By contrast, introducing heterogeneous time constants substantially improved learning performance over a homogeneous circuit with dynamical units in both the RNN and the FFL (**Fig. 8,f,g,h**). Importantly, the improvements in the learning performance with biologically inspired modifications were mirrored by an increase in the sine product, as predicted by our geometric framework (**Fig. 8i**).

Biologically inspired circuit inhibition and cellular heterogeneity not only improved learning performance at the optimal frequency but also broadened the range of frequencies that could be learned (**Fig. 8j**) To quantify this we performed Floquet analysis, which revealed that heterogeneous RNN+FFL circuits can form stable periodic attractors over a wider range of fundamental frequencies than homogeneous RNN+FFL circuits (**Fig. 8k)**.

These simulations show that incorporating biologically observed features into our simplified model of the cortico-cerebellar circuit improves time series learning performance and stability with a less effective but more biologically plausible learning rule. Our analysis of the network’s internal representations suggests that this is achieved through greater separation of task-relevant and task-irrelevant frequency components.

## Discussion

Skilled volitional movements depend on the autonomous generation of precisely timed motor commands^3,10,13^, but how interconnected motor circuits learn and stably sustain such signals remains unclear. To study time series learning in the cortico-cerebellar system we paired a cortex-like RNN with a cerebellar-like feed-forward layer in a closed-loop and trained it to generate continuous periodic signals, as required for different rhythmic behavioural tasks. The feed-forward layer enabled learning of complex periodic time series that parameter-matched RNNs could not generate. Floquet analysis revealed that the feed-forward layer enabled the formation of stable periodic attractors that were unstable in RNNs. Decomposing population activity into task-relevant and task-irrelevant frequency modes revealed that the feed-forward layer improves learning by increasing the angular separation between these subspaces and within the task-relevant subspace. Metrics of this separation and of task-relevant signal amplitude predicted learning performance and closed-loop stability across linear and nonlinear circuits, activity levels and biologically inspired heterogeneity. These results suggest that the cerebellum acts as a frequency separator that stabilises cortical dynamics during time series learning.

### Frequency separation stabilises closed-loop dynamics

Counterintuitively, the addition of a feed-forward layer improves learning and stability despite generating more task-irrelevant activity than the RNN circuit alone. This activity arises from nonlinear granule-cell-like units in the feed-forward layer introducing unwanted harmonics and phase-shifted components. Fourier analysis resolves this apparent paradox by showing that recoding in the feed-forward layer separates unwanted task-irrelevant activity from task-relevant frequency components by increasing the principal angle between their subspaces. Increased separation is due to feed-forward units acquiring more varied phase combinations than RNN units, which maintain comparatively rigid phase relationships across frequencies^25^. This enables learning rules to extract task-relevant components and discard task-irrelevant ones and prevents task-irrelevant components from being fed back into the loop, and destabilising the periodic attractor. These results extend the concept of output-potent and output-null neural subspaces^56–58^, by showing how they can be implemented in the frequency domain by feedforward circuits for rhythmic behaviours. Previous models have shown that cerebellar feedback could accelerate cortical learning^41,59^ and ease temporal credit assignment^39,40^. Our results add a distinct, geometric mechanism and provide a framework for understanding how the cerebellum supports neocortical attractor dynamics during motor tasks^21,29,30,33,34^.

### Dense codes and Golgi cell inhibition trade-off frequency separation and signal amplitude

Our findings provide new insights into why granule cell activity levels are considerably higher than the sparse code predicted by Marr-Albus theory^18,26,27,60,61^. Sparser activity increases the angular separation of task-relevant and task-irrelevant subspaces but reduces task-relevant amplitudes, a trade-off that makes learning performance optimal at intermediate activity levels. This extends theoretical work showing that learning in cerebellum-like circuits is optimal at intermediate coding levels^62–64^ to autonomous time series learning in cortico-cerebellar loops. Our results also show that cellular heterogeneity and Golgi cell feedforward inhibition of granule cells^65^, each improve performance and broaden the range of learnable frequencies. Moreover, our sine product metric revealed that both of these act by improving angular separation of task-relevant and task-irrelevant frequency components.

### Cerebellar computation depends on input properties

Marr-Albus theory proposes that the cerebellar granule cell layer performs pattern separation by nonlinearly mixing diverse sensorimotor signals and projecting them into higher-dimensional space where their sparse representations are linearly separable^26,27^. In this view of the cerebellum as a pattern classifier^26,27,66^, the embedding dimensionality of the representation predicts the associative learning performance^48^. Such pattern separation is most relevant for distinguishing between diverse contexts and different behaviours^55,61^ that together span a large number of dimensions.

By contrast, individual behavioural tasks and their neural representations are often semi-continuous and low dimensional^10,35,67^. Separation and classification of discrete patterns may not be the primary transformation required for learning such correlated neural representations. Adaptive filter models^51,68^, which consider granule cells as linear filters that provide a temporal basis for Purkinje cells to learn time varying signals, offer an alternative framework for understanding the processing and learning of low dimensional representations. Our results build on this idea by showing that the cerebellar input layer improves learning of low-dimensional periodic time series by increasing the angular separation of task-relevant and task-irrelevant components, and the amplitude of the task-relevant components. Recordings from mossy fibres and Purkinje cells in monkey oculomotor vermis during a saccade task have revealed limit cycle-like manifolds of neuronal activity^69^, consistent with our prediction of stable periodic attractors in cortico-cerebellar loops. However, the recent finding that neural manifolds undergo rotation between motor cortex and the cerebellar granule cell layer^38^ differs from our results in that it involves rotation of entire behavioural manifolds in state space rather than task-relevant and task-irrelevant frequency components in Fourier-coefficient space. Thus, rather than manifold separation, our findings reveal the precise geometric transformations involved in learning task-relevant periodic components in the cortico-cerebellar loop.

Our results suggest the cerebellar input layer circuit supports at least two distinct computations. To distinguish diverse contexts and behaviours it performs combinatorial mixing and separates activity patterns by increasing the embedding dimensionality of the representation^28^, while for low-dimensional periodic inputs it performs a geometric transformation that separates task-relevant and task-irrelevant frequency components. This dual role should help reconcile the conflicting views on cerebellar contributions to behaviour.

### A task-aligned framework for analysing recurrent circuits

Population activity is commonly analysed with variance-based methods such as PCA^49,61,70^, which have revealed high-dimensional representations of spontaneous behaviour in neocortex^70^ and granule cells^61^. Embedding dimensionality and condition number calculated with PCA predict classification performance^28,48,71,72^ and the learning of single sinusoids by linear RNNs^25^. We reproduced the latter result, but the relationship between dimensionality and learning performance was less clear when multiple frequency components were present. This is because PCA’s task-agnostic modes mix task-relevant and task-irrelevant frequency components, obscuring the geometry of the neural representation that governs learning.

Our task-informed frequency decomposition (see also ref.^73^) addresses this problem, and when combined with our sine product and component norm metrics, and Floquet analysis, provides a general framework for linking the geometry of neural representations to learning and stability for rhythmic behaviours. This is most readily applied to simple well-defined rhythmic behaviours where the task-relevant frequency components to be learned can be determined directly or from the muscles involved^35^. However, for more complex behaviours such as locomotion, identifying the task-relevant signal components to be learned by each Purkinje cell may also require the feedback error they receive, which can be read out from their complex spikes. Our framework could be applied to other recurrent loops that generate rhythmic or sequential activity, including cortico-basal ganglia-thalamic loops, hippocampal–entorhinal circuits, brainstem and spinal central pattern generators, and the song system of songbirds. In each case, it could be used to quantify the geometry of the neural representation and whether the angular separation in frequency components predicts learning performance and the stability of periodic attractors.

### Predictions

Our models make several testable predictions in addition to the presence of periodic attractor states already observed in monkey cerebellum^69^. First, during periodic behaviours such as cyclical forelimb^35^ and complex locomotion tasks^74^, the angle between task-relevant and task-irrelevant activity modes should be larger in granule cells than in pontine mossy fibres when assayed with the sine product. Related to this property, heterogeneous short-term plasticity at mossy fibre synapses^75–77^ is expected to contribute to this angular separation. Two existing observations are consistent with frequency separation occurring in the granule cell layer: the frequency response properties of cerebellar granule cells have been found to vary in a graded manner^78^ and activity in deep cerebellar nuclei precisely encodes the frequency information in periodic behaviours^37^. The second prediction is that Golgi cell inhibition should shift the sine product and norm metrics in opposite directions. Lastly, reducing frequency separation in the cerebellum should destabilise periodic attractors in the motor cortex, particularly for learned behaviours with multiple frequency components.

### Limitations

For tractability and interpretability, our models simplify the cortico-cerebellar circuit, omit feedback delays and use non-biological learning rules. While highly simplified, the rate-based RNN and feed-forward layer perceptron models we employed have been highly effective at modelling measured neocortical^11^ and cerebellar^64,76^ neural population activity. However, treating the pontine nuclei, deep cerebellar nuclei and thalamus as relays neglects the processing they perform^79^. Omitting delays may seem problematic, because delays can introduce phase shifts and destabilise closed loops^9,80^. However, since online learning rules can account for feedback delays^22^, we assumed delays introduced by the cortico-cerebellar loop are compensated for.

We used two learning rules that differ markedly in their capacity to find solutions^81^: least-squares learning and a biologically plausible covariance rule^50^ that captures some aspects of cerebellar supervised learning^51^. Obtaining consistent results with both suggest that our conclusions generalise across learning rules.

Here we have focused on time series learning of signals with multiple periodic components, but our approach could be extended to aperiodic movements. This would require other basis sets that could be derived from the behavioural data^82^. Behaviour-specific basis sets could be then used to decompose neural recordings into signal and noise subspaces. Spiking, distributed plasticity^83^ and synaptic dynamics^75–77^ also remain to be incorporated into our geometric framework. The principles linking neural representation to closed-loop learning we have identified are nonetheless likely to hold, because linear networks reproduced the stability properties of nonlinear networks once their representational geometry was matched.

## Conclusion

In summary, by linking the geometry of neural representations to closed-loop learning, our results reveal how the cortico-cerebellar loop generates and stably maintains attractor dynamics for movements with multiple periodic components. They also provide a computational rationale for combining a cerebellar feed-forward layer with recurrent cortical circuitry.

## Methods

### Cortical network model

The cortical circuit was modelled as a recurrent neural network (RNN) with N_ctx_ units and state vector 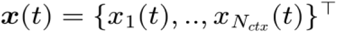 , evolving according the state equations:

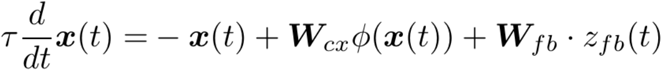

where the single neuron time-constant is τ = 10*ms*. z_fb_(t) denotes a continuous input/feedback-signal (see below) and *ϕ* (*x*) = tanh(*x*) is the neuronal activation function, applied element-wise. The recurrent weights were fixed and drawn from a zero-mean Gaussian distribution with standard deviation 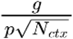 with connection probability p = 0.1 and g = 0.9 (unless stated otherwise). The feedback weights were fixed and drawn from a standard Gaussian distribution.

### Feed-forward layer model

The cerebellar-like feed-forward layer was modelled as a feedforward network with *N*_*gc*_ granule cell-like units and state vector ***y***(*t*) = {*y*_1_(*t*), .. , *y N*_*gc*_ (*t*)^⊤^ . Inputs and firing rates of instantaneous feed-forward layer units were given by:

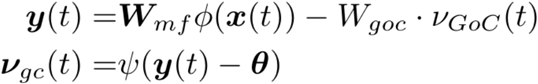

Here, W_goc_ = 0.75 represents a weight that sets the strength of feed-forward Golgi cell inhibition. We assumed Golgi cells to operate in the fluctuation driven regime^42,43^, providing global inhibition proportional to the standard-deviation of the cortical activity according to *v* _*GoC*_(*t*) = std(*x*(*t*)). Mossy fibres were assumed to faithfully relay cortical activity and were not explicitly modelled. Each feed-forward layer unit directly received exactly 4 distinct, randomly chosen inputs from the cortical units, i.e. each row of **W**_*mf*_ has exactly 4 non-zero entries. The MF weights were fixed to unity throughout all simulations. The feed-forward layer unit non-linear activation function *ψ* was chosen to be a differential variant of the linear rectifier to reflect the threshold properties of granule cells^84,85^ and was applied element-wise, with 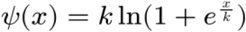, and k = 20. *θ* denotes the vector of individual granule cell thresholds, which were set as described below.

### Feed-forward layer unit thresholds

Feed-forward layer unit thresholds were set individually by driving the cortical network with the target signal for 15 periods until it reached its asymptotic periodic state, after which the activity over two periods was collected. For each feed-forward layer unit, its threshold *θ*_*i*_ was set to the corresponding quantile of its input distribution such that the net input exceeded *θ*_*i*_ for a fixed fraction of time points (i.e. 10%, 20%, etc.) during the period.

### Feedback

The input/feedback-signal received by the cortical circuit is given by:

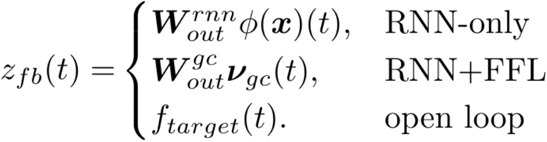

In the closed-loop case the outputs were produced by a linear readout unit connected to the RNN or the feed-forward layer, with readout weights 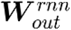 and 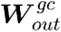 , respectively. In the open-loop case the network was driven by the target signal.

### Target signals

We used two periodic target signals, tagged as ‘simple’ and ‘complex’, and given by:

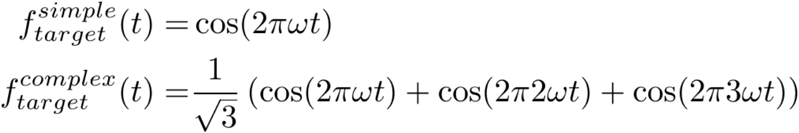

The normalisation factor ensures that both target signals have the same variance.

### Network models with dynamical units

For the networks in **Fig. 8** we introduced heterogeneous single-unit time-constants and injected white noise into the RNN network during the test-phase. RNN unit activity was now governed by the state equations:

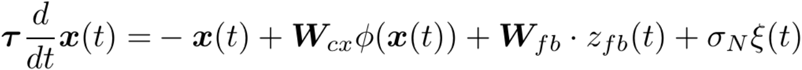

where, *ξ*(*t*) is a Gaussian white noise process with zero mean and unit variance and σ_*N*_ = 10^−3^. In **Fig. 8b** and **8g**, the single neuron time-constants τ were drawn from an exponential distribution with average τ_*avrg*_ = 10*ms*, truncated below a minimal allowed time-constant of τ_*min*_ = 3*ms*. For the networks in **Fig. 8f-g** we modelled feed-forward layer units using first-order differential equations, governed by:

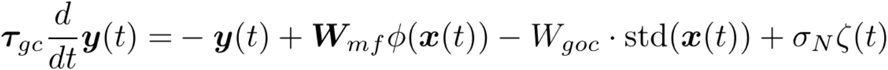

increasing the total size of the dynamical system from N_ctx_ to N_ctx_ + N_gc_. *ζ*(*t*) denotes a noise process injected during the test-phase similarly as above. Feed-forward layer unit time-constants were set to electrophysiologically determined values of granule cell membrane time-constants^52^, with τ_*gc*_ = 3.4*ms* for the network in **Fig. 8f**. For the network in **Fig. 8g**, time-constants were drawn from a gamma distribution, estimated from the same data, with average τ_*gc,avrg*_ = 3.4*ms*, standard deviation τ_*gc,std*_ = 1.2*ms* and truncated below the minimal experimental value of τ_*gc,min*_ = 1.4*ms*. We used N_gc_ = 300 in **Fig. 8a-d** and N_gc_ = 1200 in **Fig. 8e-f** and N_ctx_ = 300 in all simulations. Feed-forward layer activity levels were set to 0.4 and W_goc_ = 0 in **Fig. 8c**.

### Least-squares training

For least square learning, training was carried out as in ref^25^. The network was driven by the target signal in the open loop setup. After a transient phase of 15 target signal periods, the network was assumed to have reached its asymptotic periodic state. Network activity was then sampled at a time step of 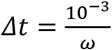 and collected over 10 target signal periods into the N x T matrix **X**, where N ∈ {N_ctx_, N_gc_} is the number of RNN units or feed-forward layer units and *T* is the number of time-points. Similarly, the target signal was collected into the 1 x T vector **F**_target_. We regularised network activity via 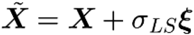 , where ξ is a random matrix of standard Gaussian variables and σ_*LS*_ = 10^−2^. Readout weights were then calculated using:

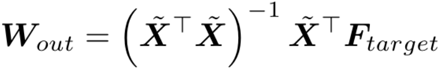

### DELTA rule training

For **Fig. 8**, training was carried out in the closed-loop setup, using a delta rule variant of FORCE learning^22^. The updatable readout weights were expressed as a state vector ***W***_*out*_(*t*) = {*W*_*out,1*_(*t*),…, *W*_*out,N*_*(t)*}^⊤^ with N ∈ {N_ctx_, N_gc_}, and obeyed a set of differential equations which were evolved simultaneously with the network state equations, according to:

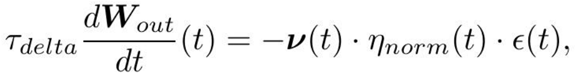

where τ_*delta*_ = 10*ms* and *v(t) ∈ {ϕ (x(t))*,***v***_*gc*_(*t*)}. The second factor represents the effective learning rate

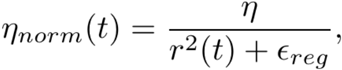

where *η* = 20, *ϵ*_*reg*_ = 10^−6^ and the normalisation 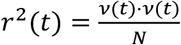 is the mean squared activity of the cortical units or feed-forward layer units at time *t*, which improves the rule’s convergence independent of the firing rate magnitudes^86^. The third factor denotes the instantaneous error at time *t* as given by:

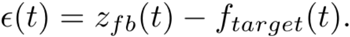

This learning rule is similar to the covariance rule^50^ that is thought to be realised by climbing fibre signalling^51^. Training of all networks in **Fig. 8** was carried out for 500 target signal periods.

### Error measure

Deviation of the output signal from the target during the closed-loop test phase was quantified by the root mean square error (RMSE):

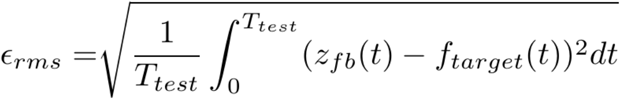

where T_test_ is the length of the test phase interval. For all simulations the test phase was set to 20 target signal periods.

### Spanning vectors, principal angles and dimensionality

Networks were driven in open-loop configuration by the target signal for 15 periods. After reaching its asymptotic periodic state, network activity was collected over 3 target signal periods. Then, the firing rate *r*_*i*_(*t*) of each individual unit was decomposed into a Fourier series according to:

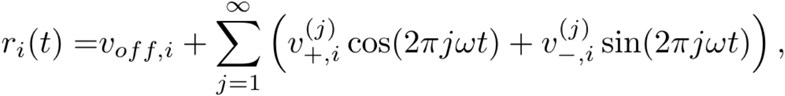

yielding the population level spanning vectors *v*_*off*_,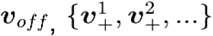 and 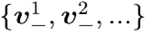 . In practice, we retained the first 60 spanning vectors for non-linear networks and a smaller number of non-zero spanning vectors for linear networks.

Spanning vectors were then centred and termed task-relevant (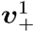 and 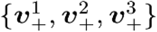 for the simple and complex target signal, respectively) or task-irrelevant (all other vectors). Principal angles between subspaces spanned by task-relevant and task-irrelevant vectors were computed from orthonormal bases using the standard singular value decomposition (SVD) approach^87^.

The sine product was computed according to:

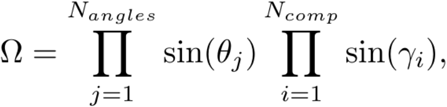

Where the γ_*i*_ are the principal angles between task-relevant and task-irrelevant subspaces of which there are as many as the number of task-relevant components, N_comp_. The θ_*i*_are the pairwise angles between task-relevant spanning vectors, their number being 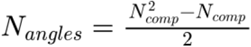 . For the simple signal (N_comp_ = 1), there are no task-relevant angles and we thus set the corresponding product to one.

In **Fig. 6d**, the dimensionality and condition number were calculated by applying principal component analysis to the network activity. The resulting eigenvalues λ_1_ ≥ λ_2_ ≥. .. of the covariance matrix were used to compute the dimensionality (D) as the participation ratio:

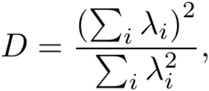

while the condition number (CN) was obtained as the ratio of the largest to the smallest non-zero eigenvalue:

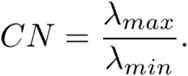

### Linear network with artificial task-irrelevant inputs

In **Fig. 5** and **Fig. 6** we used a linear version of RNN+FFL. Artificial task-irrelevant inputs can be injected either into the recurrent network or the feed-forward layer. Doing the former, however, results in additional task-irrelevant components generated by the recurrent dynamics. Thus, to be able to fully control the additional components, we chose the simpler alternative of injecting artificial inputs into the linear feed-forward layer. We employed a network with N_ctx_ = N_gc_ = 500 , g = 0.75 and state vectors **x**(t) and **y**(t) as above, evolving according to

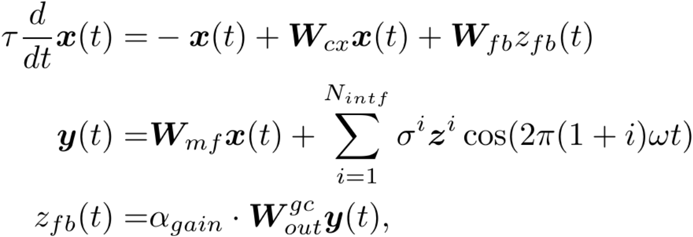

where α_*gain*_ is a global feed-forward layer unit gain. The network was trained via least squares to produce the simple target signal 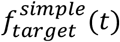 with ω = 10Hz. Feed-forward layer activity was driven, firstly, by mossy fibre inputs from the cortical network, ***W***_*mf*_***x***(*t*), featuring two spanning vector components *v*_+_ and *v*_−_ ; secondly, by N_intf_ periodic task-irrelevant inputs that were injected into the feed-forward layer with unit-norm spanning vectors *z*^*i*^ with zero-mean Gaussian elements. Angles between task-relevant and task-irrelevant components are described by the correlation matrix:

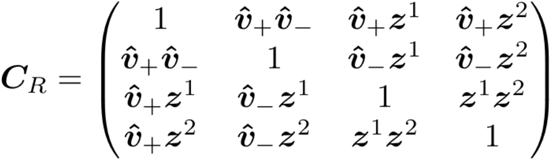

where 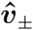 denotes the normalised spanning vectors,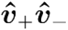 was fixed by the recurrent cortical dynamics and 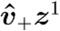 and 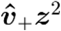 were treated as fixed parameters. Modification of 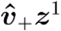 and/or 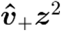 potentially led to correlation matrices that were not positive semidefinite due to the combination of free and fixed entries. To obtain the nearest valid correlation matrix while preserving the fixed parameters, we adjusted all non-fixed entries by applying alternating projections between the positive semidefinite cone and the constraint set^88^, with initial 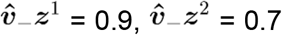 and *z*^1^ *z*^2^= 0.8. The adjustment was terminated when the smallest eigenvalue exceeded -10^-8^ or after 1000 iterations.

We used the following additional parameters. Fig. 5a-e: N_intf_ = 1, α_*gain*_ = σ^1^ = 1 while scanning over 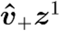 . Fig. 5f-j: N_intf_ = 2, α_*gain*_ = σ^1^ = σ^2^ =1, 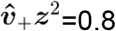 while scanning over 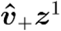 . Fig. 5k-o: N_intf_ = 2, α_*gain*_ = σ^1^ = σ^2^ =1, 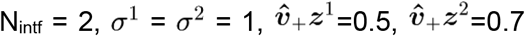 while scanning over α_*gain*_ . Fig. 6: N_intf_ = 1, α_*gain*_ = 1, 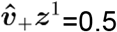 while scanning over σ^1^ .

### Computation of largest Floquet multiplier

To assess the stability of the target periodic attractor, or limit cycle, we computed the largest non-trivial Floquet multiplier^44,45^ using a single perturbation vector evolved under linearised dynamics. We defined an augmented system, consisting of the state equations above and the variational equation

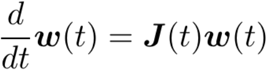

where **w**(t) is a random, N dimensional perturbation vector and **J**(t) is the RNN’s or RNN+FFL’s Jacobian. The size of the augmented system was thus 2N_ctx_ for networks with instantaneous feed-forward layer units and 2(N_ctx_ + N_gc_) for the network in **Fig. 8g**.

The state equations and the variational equations were treated asymmetrically: the state equations were evolved in open-loop, i.e. were driven by the external target signal throughout the Floquet integration, keeping the system’s trajectory on the target limit cycle ***x***^*^(*t*). The Jacobian **J**(t), on the other hand, was evaluated using the closed-loop feedback term arising from the learned readout weights **W**_out_. The explicit forms of **J**(t) for each network architecture (RNN only, or RNN+FFL with instantaneous or dynamic feed-forward layer units) are provided below. Importantly, our asymmetric approach addresses the question whether the target periodic attractor is a stable solution of the closed-loop system after learning. This is distinct from the alternative fully closed-loop Floquet analysis, which assesses stability of whatever closed-loop limit cycle emerges after learning rather than the target limit cycle. Our approach instead asks whether, given that the network is on the target limit cycle, the closed-loop feedback learned during training acts to stabilise or destabilise it.

To carry out the computation, networks were driven in open-loop configuration by the target signal for 15 periods. After reaching its asymptotic periodic attractor state the augmented system was initialised from the network state at the end of this transient. An initial perturbation vector **w**_0_ was drawn from a standard Gaussian distribution and normalised to unit norm and Floquet integration was carried out over K = 10 periods. At the end of each period, the trivial Floquet direction, corresponding to perturbations along the limit cycle, was projected out by computing

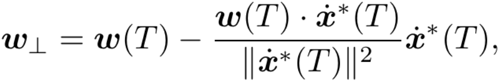

where T is the length of the period and 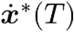 is the tangent vector to the target trajectory at the end of the period. The log norm log(||**w**_⊥_||) was accumulated and **w**_⊥_ was renormalised to unit norm before integration of the next period. The largest non-trivial Floquet multiplier was then computed according to:

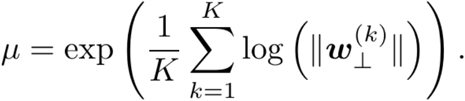

multiplier μ < 1 indicates a stable periodic attractor and μ > 1 instability. The number of integration periods (K = 10) was chosen empirically: the resulting multipliers were consistent with the observed network phenomenology across all conditions and architecture, suggesting sufficient convergence.

In **Fig. 1, Fig. 7, Fig. 8** and **Fig. S1** parameter configurations were considered unstable when ≥20% of simulations had multipliers larger than unity. The networks employed in **Fig. 5** were overall more stable due to their entirely linear dynamics. We therefore used a lower cutoff of ≥10% for this figure.

### Jacobians for Floquet computation

The form of the Jacobian **J**(t) depends on the network architecture that is integrated. Here we list the Jacobians for the non-linear RNN-only and RNN+FFL with and without dynamic feed-forward layer units.

Suppressing the time argument for readability, the RNN-only Jacobian is an N_ctx_ x N_ctx_ matrix of the form:

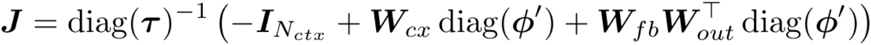

where 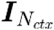 is the N_ctx_ x N_ctx_ identity matrix and ϕ′ is evaluated along the driven limit cycle trajectory *x*^*^(*t*) and τ is the vector of (homogeneous or heterogeneous) single unit time constants.

The Jacobian for RNN+FFL with instantaneous feed-forward layer units is likewise an N_ctx_ x N_ctx_ matrix:

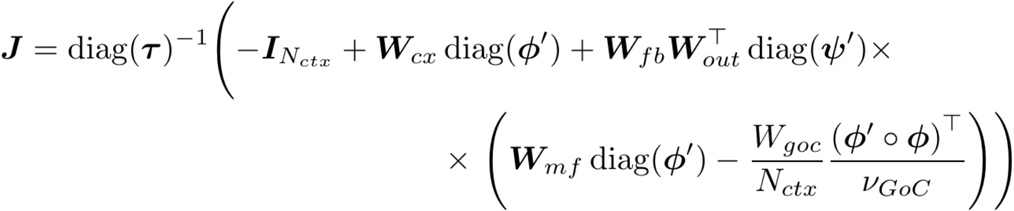

where similarly ϕ, ψ′ and *v*_*GoC*_ are evaluated along ***x***^*^(*t*).

Finally, for the RNN+FFL with dynamic feed-forward layer units the Jacobian increases to an (N_ctx_ + N_gc_) x (N_ctx_ + N_gc_) matrix:

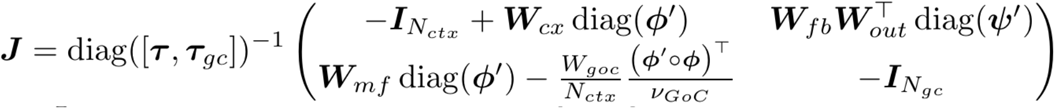

where 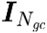 is the (N_ctx_ + N_gc_) identity matrix and [τ, τ_*gc*_] represents the concatenation of the time constant vectors.

The cases with linear activation functions and zero Golgi cell inhibition are special cases of the above.

### Amplitude of readout weights for activity with two task-relevant components and one task-irrelevant-component

To generate the plots in **Fig. S2**, we analytically calculate the amplitude (norm) of the least-squares readout weights for a linear network with a task-relevant space spanned by two task-relevant component vectors, *s*_1_ and *s*_2_, and a task-irrelevant space spanned by one task-irrelevant component vector, *n*. While this abstract case cannot be realised in a simulation, it provides insights into the dependence of ║***W***_*out*_ ║ on the angle θ between the task-relevant vectors for a given principal angle γ between task-relevant- and task-irrelevant-subspaces. In the following, we assume for the sake of simplicity that the angles between the task-irrelevant component and each of the task-relevant components are equal, i.e. ϕ = ∠ (s_1_ , *n*) = ∠ (s_1_ , *n*). We first provide an expression for the principal angle between the task-relevant- and task-irrelevant-subspaces. In a second step, we then derive a formula for ║***W***_*out*_║.

Let N denote the task-irrelevant-subspace, spanned by the unit norm vector 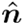 and S denote the task-relevant-subspace spanned by the unit norm vectors 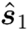 and 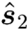 . Using the Gram-Schmidt procedure, we can find 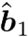 and 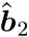, the orthonormal basis vectors spanning S. They read:

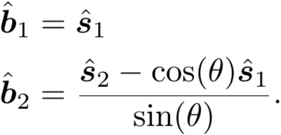

The principal angle between S and N is equal to the angle between 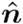 and its normalised projection onto S:

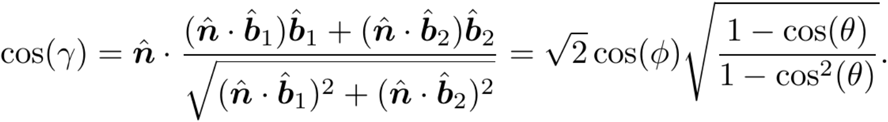

Thus, expressed in function of the principal angle γ is given by:

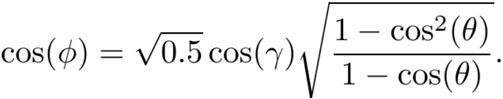

Following ref.^25^, the least-squares solution is determined by following condition:

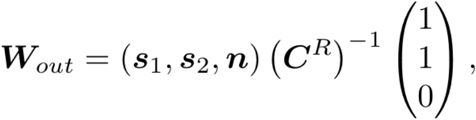

where (*s*_1_, *s*_2_,*n*) is the collection of the spanning vectors of the network activity and the column vector (1,1,0)^*t*^ denotes the Fourier-space representation of the target signal. The matrix *C*_*R*_ is the reduced 3×3 correlation matrix of the network activity given by:

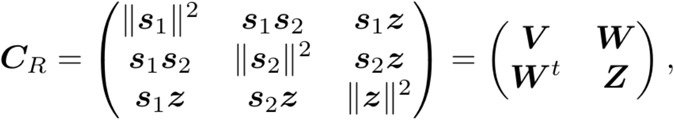

Where ***V*** and ***W*** are the corresponding 2×2 and 2×1 block-matrices, respectively and ***Z =*** ║ *z ║*^2^ . The inverse of *C*_*R*_ is given by:

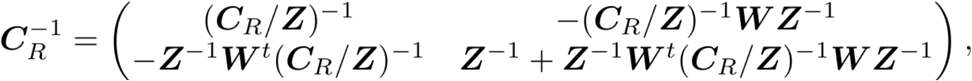

where the symbol *C*_*R*_*/Z* denotes the Schur complement with respect to block *Z*:

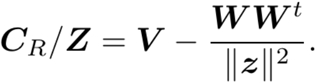

The inverse of the Schur complement is:

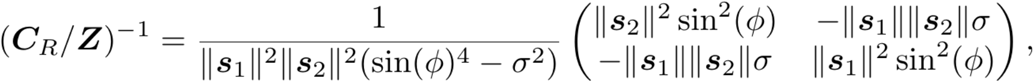

where σ = cos(θ) −. cos^2^(ϕ).

Inserting the above and carrying out the algebraic manipulations we obtain:

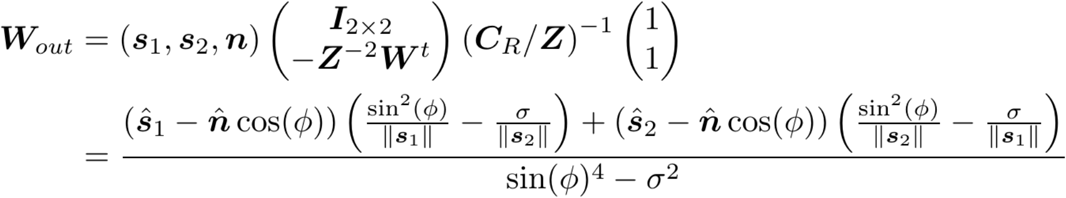

With this, we can derive ║**W**_*out*_║ :

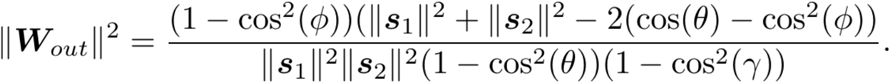

Together with the equation for ϕ ║**W**_*out*_║, can be computed as a function of the norms ║*s*_1_║, ║*s*_2_║ and the angles θand γ.

### Numerical integration

Most simulations, including the computation of the Floquet multiplier, were carried out using the Runge-Kutta-Fehlberg(4,5) method with adaptive step-size control provided by the Gnu Scientific Library^89^ with absolute and relative error bounds of 10^-3^. Where extraction of the population or single unit activity was necessary, network activity was sampled with a time step 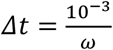. In the simulations for **Fig. 8**, the injection of white noise into the networks during the test phase made Runge-Kutta integration unfeasible. In these cases, we carried out numerical integration with the Euler-Mayurama method and a step size of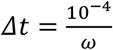. For every parameter set, results were averaged over 50 simulations with independent realisations of the connectivity matrices and noise processes.

### Statistics and reproducibility

As this is a theoretical study no experimental data was produced. Simulations were carried out with fixed random number seeds, which will be provided with the code and the scripts to generate the figures upon publication.

## Supplementary material

**Figure S1.**
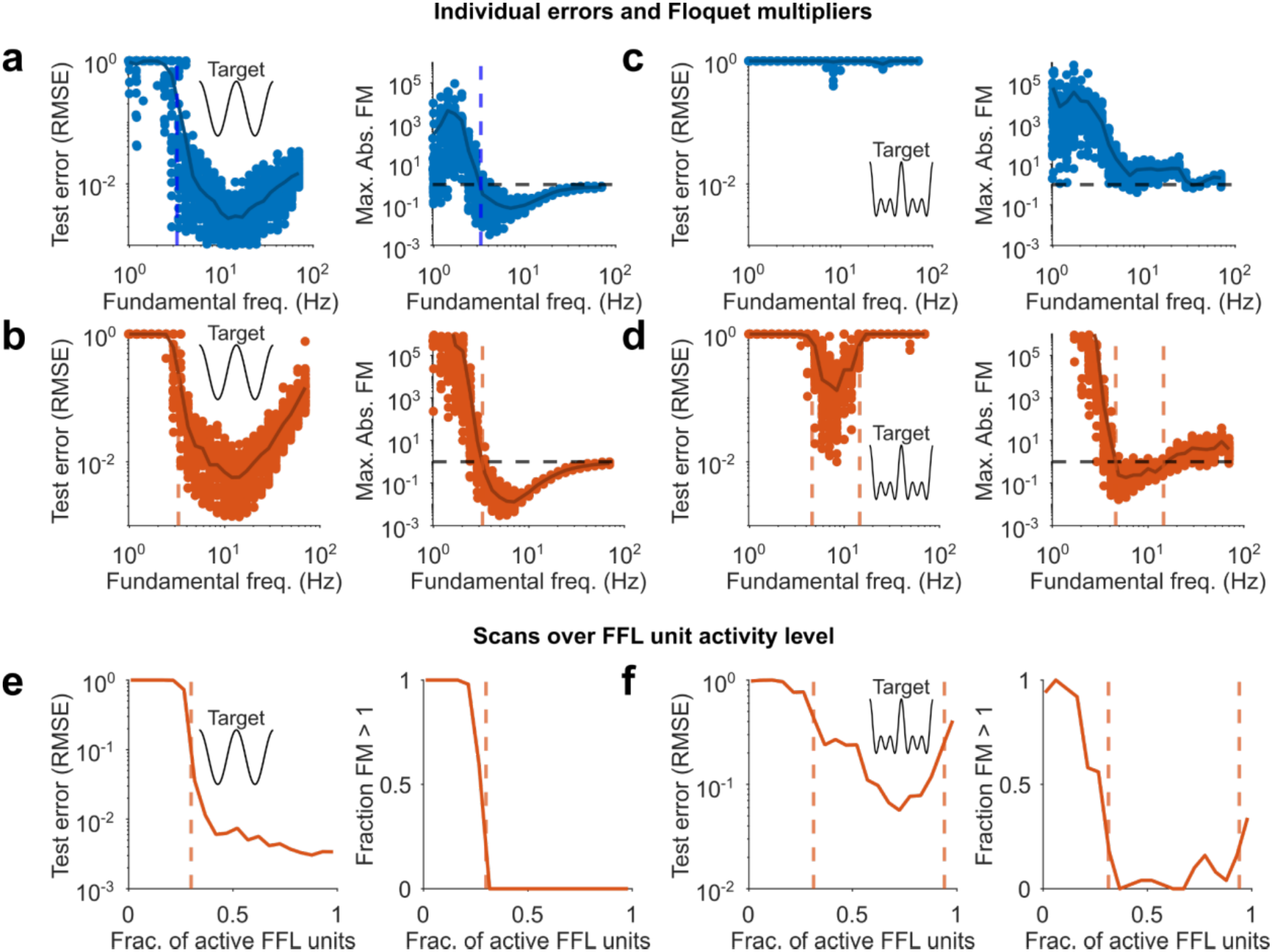
Individual errors and Floquet multipliers of the curves shown in Fig. 1 and scans over the feed-forward layer unit activity level. **a** Individual root mean square error (RMSE; left) and individual largest non-trivial Floquet multipliers for RNN-only trained on the simple signal; dark lines are averages. **b** As (**a**) but for RNN+FFL. **c**,**d** As (**a**,**b**) but for complex signal. **e** Simple signal training. Left: normalised RMSE versus feed-forward layer unit activity level; dashed red line: stability margin obtained from largest non-trivial Floquet multipliers; the target’s fundamental frequency is 10Hz. Right: fraction of largest non-trivial absolute Floquet multipliers larger than 1 versus feed-forward layer unit activity level. **f** Same as (**e**), but for complex signal.

**Figure S2.**
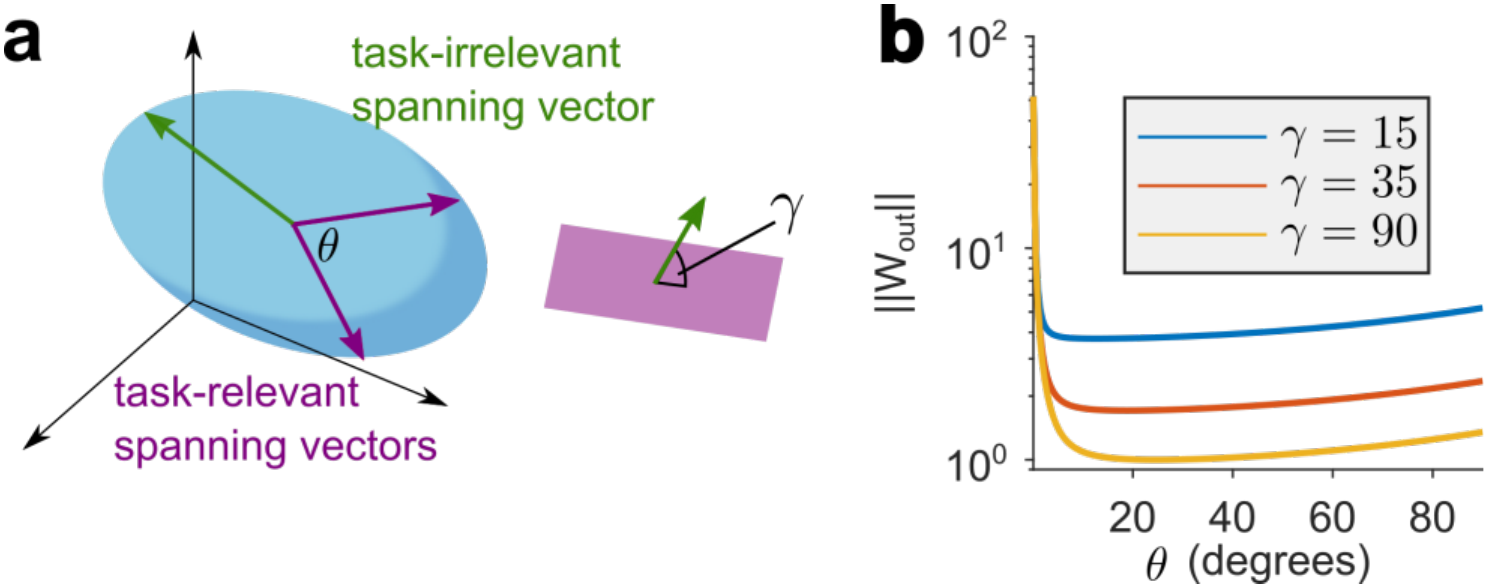
The impact of the angle between task-relevant spanning vectors on the norm of the readout weights. **a** Hypothetical population activity composed of two task-relevant spanning vectors (magenta arrows) and one task-irrelevant spanning vector (green arrow); θ : angle between task-relevant vectors; γ : principal angle between task-relevant and task-irrelevant subspaces. **b** Norm of readout weights as a function of θ for various values of γ calculated analytically; the ratio of the norms of the task-relevants spanning vectors is 1.1 (see **Methods**).

**Figure S3.**
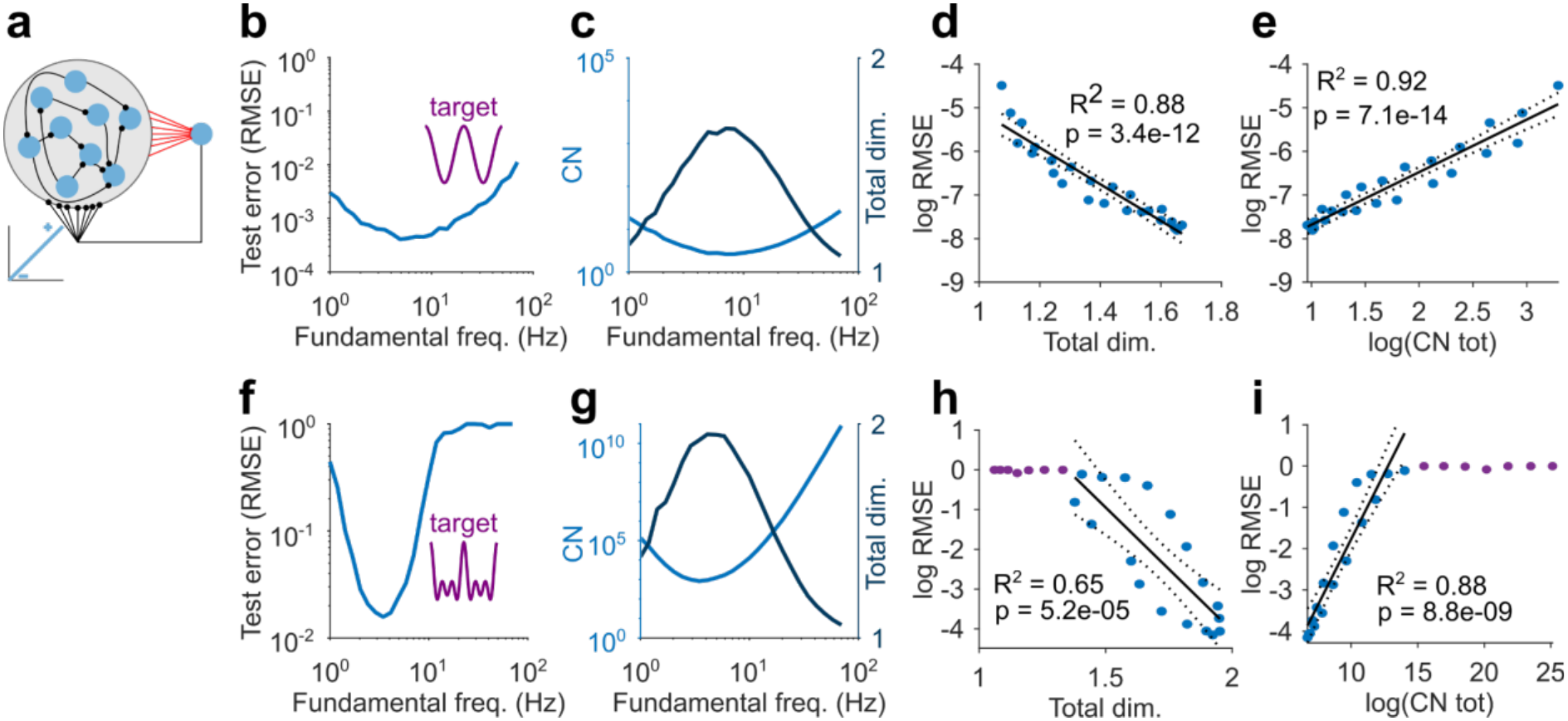
Predicting test-phase error with dimensionality and condition number in linear RNNs. **a** Schematic of linear RNN. **b** Normalised root mean square error (RMSE) versus target signal fundamental frequency for simple signal; lines represent averages over 50 simulations. **c** Condition number (CN, blue line) and total dimensionality (black line) versus fundamental frequency. **d** Logarithmic RMSE versus total dimensionality; same simulated data as in (**b** and **c**). Solid and dotted black lines indicate linear fit and 95% confidence bounds, respectively. **e** Same as (**d**) but for logarithmic condition number. **f-g** Same as (**b-e**), but for complex target signal. The magenta points in (**h**) and (**i**) indicate simulated data points with RMSE>0.9; these were excluded from the linear regression to remove biases due to the observed saturation of the RMSE close to unity.

### Data availability

Simulation results in the main figures and in extended data figures will be made available on publication on GitHub https://github.com/SilverLabUCL/Cortico-Cerebellar-Model-public

### Code availability

The models and analysis scripts will be made available on publication on GitHub https://github.com/SilverLabUCL/Cortico-Cerebellar-Model-public

## Acknowledgements

This project was supported by the Wellcome Trust (203048; 224499). R.A.S. is in receipt of a Wellcome Trust Principal Research Fellowship (224499). A.B. thanks Martin T. Wiechert for insightful discussions. We thank Sadra Sadeh and Lavinia Mitiko Takarabe for their comments on the manuscript. Anthropic’s Claude was used as an aid in the editing process and to check the code. It was also used to generate sections of the code (calculation of the correlation matrix for artificial inputs and for the accumulation of the largest non-trivial Floquet multiplier over multiple cycles).

## Author contributions

Conceptualization: A.B., H.G. and R.A.S. Methodology: A.B. and R.A.S. Software: A.B. Formal analysis: A.B. Investigation: A.B. and R.A.S. Writing—original draft preparation: A.B and R.A.S. Writing—review and editing: A.B. and R.A.S. Supervision: R.A.S. Funding acquisition: R.A.S.

### Competing interests

The authors declare no competing interests.

## Notes

### Competing Interest Statement

The authors have declared no competing interest.

